# Controlled retrieval relies on directed interactions between semantic control regions and visual cortex: MEG evidence from oscillatory dynamics

**DOI:** 10.1101/2024.09.07.611827

**Authors:** Susanne Eisenhauer, Meichao Zhang, Katya Krieger-Redwood, Richard Aveyard, Rebecca L. Jackson, Piers L. Cornelissen, Jonathan Smallwood, Elizabeth Jefferies

**Affiliations:** Department of Psychology, University of York, YO10 5DD, UK; York Neuroimaging Centre, Innovation Way, York, YO10 5DD, UK; Basque Center on Cognition, Brain and Language, 20009, Donostia-San Sebastián, Spain; CAS Key Laboratory of Behavioural Science, Institute of Psychology, Chinese Academy of Sciences, Beijing 100101, China; Department of Psychology, University of Chinese Academy of Sciences, Beijing 100049, China; Department of Psychology, Northumbria University, NE1 8ST, Newcastle upon Tyne, UK; Department of Psychology, Queen’s University, K7L 3N6, Kingston, ON, Canada

## Abstract

To navigate the world, we store knowledge about relationships between concepts and retrieve this information flexibly to suit our goals. The semantic control network, comprising left inferior frontal gyrus (IFG) and posterior middle temporal gyrus (pMTG), is thought to orchestrate this flexible retrieval by modulating sensory inputs. However, interactions between semantic control and input regions are not sufficiently understood. Moreover, pMTG’s well-formed structural connections to IFG and visual cortex suggest it as a candidate region to integrate control and input processes. We used magnetoencephalography to investigate oscillatory dynamics during semantic decisions to pairs of words, when participants (both sexes) did or did not know the type of semantic relation between them. IFG showed increases and decreases in oscillatory activity to prior task knowledge, while pMTG only showed positive task knowledge effects. Furthermore, IFG provided sustained feedback to pMTG when task goals were known, while in the absence of goals this feedback was delayed until receiving bottom-up input from the second word. This goal-dependent feedback coincided with an earlier onset of feedforward signalling from visual cortex to pMTG, indicating rapid retrieval of task-relevant features. This pattern supports a model of semantic cognition in which pMTG integrates top-down control from IFG with bottom-up input from visual cortex to activate task-relevant semantic representations. Our findings elucidate the separate roles of anterior and posterior components of the semantic control network and reveal the spectro-temporal cascade of interactions between semantic and visual regions that underlie our ability to flexibly adapt cognition to the current goals.

**Significance Statement:** Using magnetoencephalography, we characterize the spectro-temporal dynamics that underlie our ability to flexibly adapt semantic cognition to the current context and goals. We find that semantic goals increase oscillatory activity in IFG and pMTG, and ultimately facilitate visual processing. Effective connectivity analyses reveal more sustained feedback from IFG to pMTG, and more rapid feedforward signalling from visual cortex to pMTG, resulting in rapid retrieval when semantic goals are known. Crucially, our findings suggest differential roles for the two semantic control regions: while IFG *controls* goal-dependent retrieval, pMTG *integrates* top-down information from IFG with bottom-up visual input.

## 1 Introduction

Semantic memory is thought to draw on multimodal abstraction across experiences to extract knowledge about objects, words, and their relations, which we flexibly retrieve to suit the circumstances (Lambon Ralph et al., 2017). For example, to prepare a fruit pie, we exploit *thematic* knowledge that apples are used as a pie filling and locate the relevant aisle in the supermarket by retrieving *taxonomic* knowledge that apples are fruit. Semantic control implemented within the left inferior frontal gyrus (IFG) and posterior middle temporal gyrus (pMTG; Noonan et al., 2013; Jackson, 2021) is thought to support this ability to flexibly adapt semantic retrieval to suit our goals (Jefferies, 2013). Previous functional magnetic resonance imaging (fMRI) studies implicated these brain regions in demanding semantic tasks, such as when two words are weakly as opposed to strongly related (Gao et al., 2021; Jackson, 2021). In contrast, fewer studies have focused on the role of these regions when the semantic context (Grindrod et al., 2008) or the goal for retrieval (Zhang et al., 2021) is known in advance, facilitating semantic cognition. Zhang et al. (2021) investigated the ability to flexibly exploit taxonomic or thematic knowledge based on our goals, using fMRI. Participants decided if two sequentially presented words were related, and on half the trials a preceding cue revealed whether the upcoming words would share a taxonomic or thematic relation. Knowledge of the relation increased activation in IFG, indicating its role in goal-dependent semantic cognition. In addition, an interaction between this task knowledge and the presentation order of the words emerged in visual cortex: activation was stronger to the first word when the retrieval goal was known, and stronger to the second word when the goal was unknown (Zhang et al., 2021). This suggests that semantic goals enable an earlier onset of visual processing, potentially reflecting early retrieval of features that are relevant to process the anticipated relation between the two words, while without prior knowledge feature retrieval relies more heavily on the second word.

However, the relative timing and directed interactions of these goal-dependent effects in IFG and visual cortex cannot easily be inferred from fMRI findings. Consequently, it remains unclear whether regions supporting semantic control play a direct, early role in the gating of visual activation during goal-dependent cognition (Jackson et al., 2021). While connections from frontal to occipital cortex support word reading (Schurz et al., 2014; Woodhead et al., 2014), it is unclear whether prior semantic knowledge modifies such connections. Furthermore, computational modelling indicated the need of a region that integrates bottom-up input with controlled top-down processes in semantic cognition (Giallanza et al. 2024). pMTG is a candidate for such a process since it is well connected anatomically to both visual cortex and IFG (Turken and Dronkers, 2011). Directional time-resolved connectivity measures can address these issues and confirm whether pMTG receives both top-down input from IFG and bottom-up input from visual cortex.

Here, we used magnetoencephalography (MEG) to investigate the spectro-temporal dynamics and directional connectivity between semantic control regions and visual cortex during goal-oriented semantic cognition. We used Zhang et al.’s paradigm (2021), in which participants either knew or did not know whether a subsequent semantic judgement would be based on a taxonomic or thematic link. In time-frequency space, we asked whether effects of task knowledge in semantic control regions (IFG and pMTG) precede interactions of task knowledge with bottom-up input in visual cortex (Fig. 1a). We then used spectral Granger Causality to investigate the effective connectivity between semantic control regions and visual cortex. If interactions between task goals and visual processes are driven by semantic control regions, top-down effects should precede or coincide with bottom-up inputs (Fig. 1b). Finally, we delineated the role of pMTG by investigating whether it provides top-down constraint over visual cortex or integrates top-down influences from IFG with bottom-up inputs (Fig. 1c). In this way we elucidate the architecture of controlled semantic retrieval over time and frequency.

**Figure 1.**
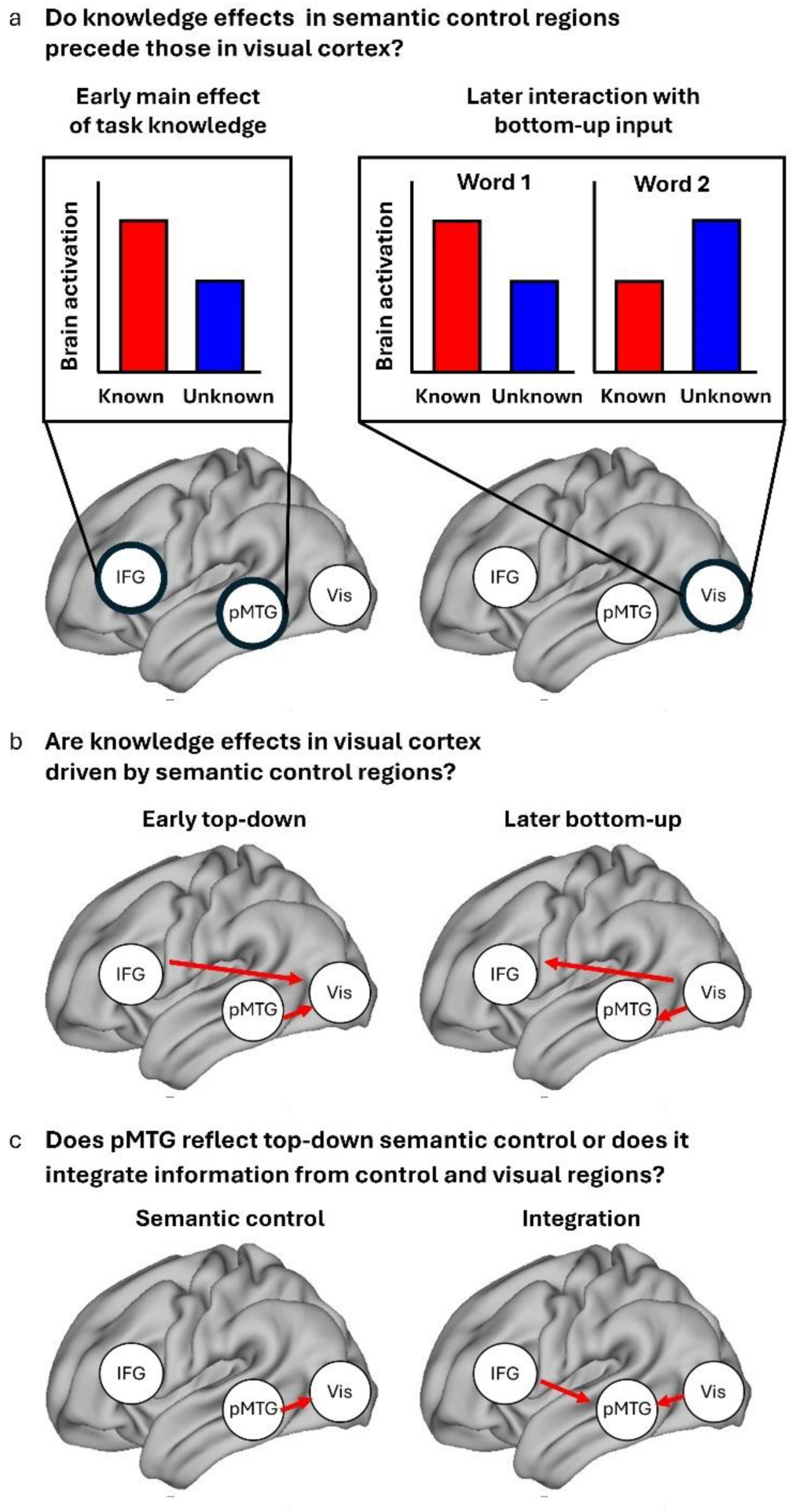
Research questions and predictions: **a)** Higher brain activation in semantic control regions (left IFG and pMTG) for known vs. unknown semantic goals (left) is expected to precede interactions between task knowledge and word order in visual cortex (right). For the interaction in visual cortex, we expect that during the presentation of the first word, brain activation is higher in the known goal condition while for the second word, activation is higher in the unknown goal condition (Zhang et al., 2021). The use of MEG allows us to specify the time windows and frequency bands underlying these effects. **b)** If task knowledge effects in visual cortex are driven by semantic control regions, we expect to observe an early top-down connection from semantic control regions to visual cortex in the known condition (left), which precedes or coincides with the bottom-up connection in the reverse direction (right). **c)** Connectivity patterns can further discriminate the functional role of left pMTG: If this region is predominantly involved in semantic control (cf. Jackson, 2021), we expect to observe top-down connections from this region to visual cortex (left). However, if pMTG (additionally) serves the integration of task knowledge and bottom-up input, it should receive inputs from both semantic control regions (IFG) and visual cortex (right).

## 2 Methods

### 2.1 Participants

#### Behavioural study

In order to investigate the effect of top-down goals on semantic decisions, a behavioural study examined the performance of 20 participants (18 females, mean age 20.7±2.9 years, range 18-29 years; 16 right-handed) who did not complete the neuroimaging investigation, using a version of the task in which responses were made on each trial. This behavioural study was included since in the MEG version, participants were instructed to only respond to unrelated catch trials, in order to prevent contamination of the related trials from motoric activation. In addition, we selected word pairs for the MEG study based on the behavioural results (see section 2.2). Participants were native English speakers and did not have any language disorders or psychiatric conditions. Of the original sample of 25 participants, five were excluded due to insufficient task performance (i.e., below 50% correct responses, N = 4) or not meeting the inclusion criteria (N = 1). Participants gave informed consent before participation and received a voucher or course credit as compensation. The study was approved by the ethics committee of the Department of Psychology, University of York.

#### MEG study

Twenty-eight participants (21 females, mean age 20.2 years, range 18-27 years) completed the MEG study and were entered into final data analysis. Participants were right-handed native English speakers with (corrected-to-) normal vision and did not have any language disorder or psychiatric condition. Of the original sample of 35 participants, seven were excluded due to poor task performance (N = 1, only 60% of unrelated trials detected), insufficient number of trials (<40 per condition) after artifact rejection (N = 5) and not completing the experiment (N = 1). Participants gave informed consent before participation and received vouchers or course credit as compensation. The study was approved by the ethics committee of York Neuroimaging Centre, University of York.

### 2.2 Stimulus materials

A total of 392 word pairs were presented in a behavioural semantic priming study, including 179 word pairs from Zhang et al. (2021), 84 additional word pairs from Teige et al. (2019), and 129 new word pairs. 153 word pairs were categorically related, for example, both words could be birds (peacock – penguin) or vehicles (bicycle – lorry). 179 word pairs were thematically related, i.e., the two items frequently occurred in the same context (e.g., coral – diver or armchair – fireplace). Sixty word pairs were unrelated. Word pairs were rated for thematic and taxonomic relatedness by different samples of participants (n = 30 for the materials of the original studies; Teige et al., 2019; Zhang et al., 2021; n = 27 for the new materials) on a Likert scale from 1 to 7 (1 = Not at all, 7 = Very). These ratings confirmed that categorical word pairs (mean ± standard deviation: 4.67 ± 0.92) were more visually similar than thematic word pairs (mean ± standard deviation: 1.68 ± 0.93; *t* = 29.5, *p* < 2.2e-16), while thematic word pairs (mean ± standard deviation: 4.74 ± 0.93) were rated as being found together more regularly than categorical word pairs (mean ± standard deviation: 2.92 ± 1.10; *t* = 19.6, *p* < 2.2e-16). Based on the behavioural results, we selected a subset of 120 categorically related, 120 thematically related and 60 unrelated word pairs for the MEG study which were responded to correctly by the majority of participants in the behavioural study (82.9% on average; range: 40-100%). In this subset, prime and target words as well as words from categorical and thematic pairs were matched for word length, word frequency and imageability using ‘Match’ (van Casteren and Davis, 2007). Stimulus characteristics are presented in Table 1.

**Table 1.** Stimulus Characteristics.

| **Table 1.** Stimulus Characteristics. |  |  |  |  |  |  |  |
| --- | --- | --- | --- | --- | --- | --- | --- |
|  |  | **Minimum** | **1st**<br>**Quartile** | **Median** | **Mean** | **3rd**<br>**Quartile** | **Maximum** |
| **Word 1**<br>**Length** | **Taxonomic** | 3.00 | 4.00 | 5.00 | 5.55 | 7.00 | 10.00 |
|  | **Thematic** | 3.00 | 4.00 | 6.00 | 5.77 | 7.00 | 9.00 |
|  | **Unrelated** | 3.00 | 5.00 | 5.50 | 5.78 | 7.00 | 10.00 |
| **Word 2**<br>**Length** | **Taxonomic** | 3.00 | 4.00 | 5.00 | 5.61 | 7.00 | 10.00 |
|  | **Thematic** | 3.00 | 4.00 | 5.00 | 5.65 | 7.00 | 10.00 |
|  | **Unrelated** | 2.00 | 4.00 | 5.00 | 5.75 | 7.00 | 10.00 |
| **Word 1**<br>**Frequency** | **Taxonomic** | 0.00 | 0.61 | 0.92 | 0.98 | 1.33 | 2.44 |
|  | **Thematic** | 0.00 | 0.65 | 1.02 | 1.03 | 1.36 | 2.75 |
|  | **Unrelated** | 0.14 | 0.61 | 0.96 | 0.96 | 1.27 | 2.13 |
| **Word 2**<br>**Frequency** | **Taxonomic** | 0.19 | 0.69 | 0.93 | 1.03 | 1.27 | 2.59 |
|  | **Thematic** | 0.00 | 0.80 | 1.09 | 1.10 | 1.42 | 2.10 |
|  | **Unrelated** | 0.21 | 0.56 | 0.92 | 1.02 | 1.37 | 2.44 |
| **Word 1**<br>**Imageability** | **Taxonomic** | 276.00 | 555.00 | 586.00 | 573.10 | 602.00 | 652.00 |
|  | **Thematic** | 376.00 | 541.00 | 580.00 | 564.50 | 607.50 | 655.00 |
|  | **Unrelated** | 390.00 | 553.50 | 587.00 | 575.70 | 614.80 | 640.00 |
| **Word 2**<br>**Imageability** | **Taxonomic** | 232.00 | 560.50 | 580.50 | 575.80 | 609.00 | 655.00 |
|  | **Thematic** | 232.00 | 553.50 | 581.20 | 565.90 | 605.00 | 642.00 |
|  | **Unrelated** | 398.00 | 545.80 | 595.00 | 578.30 | 612.00 | 652.00 |
| **Semantic**<br>**Similarity**<br>**(word2vec)** | **Taxonomic** | 0.10 | 0.25 | 0.32 | 0.32 | 0.40 | 0.62 |
|  | **Thematic** | 0.04 | 0.19 | 0.27 | 0.27 | 0.34 | 0.60 |
|  | **Unrelated** | -0.04 | 0.03 | 0.08 | 0.08 | 0.13 | 0.26 |

### 2.3 Experimental procedure and task

The behavioural study was carried out online using the Gorilla Experiment Builder (www.gorilla.sc; Anwyl-Irvine, et al., 2020). Words were presented in black in front of a white background (Fig. 2). The procedure followed that of Zhang et al. (2021) with shorter presentation durations. Each trial started with the presentation of two vertical black bars (nonius lines) indicating the center of the screen where participants were asked to fixate. Nonius lines were presented for a jittered duration between 800 and 1800 ms. Then, a cue which either read ‘Category?’, ‘Thematic?’, or ‘Them/Cat?’ was presented for 600 ms. In known goal trials, word pairs were from the same category or thematically related, and preceded by the respective cue (‘Category’ and ‘Thematic’, respectively). In unknown goal trials, word pairs that were from the same category or thematically related were preceded by an unspecific cue (‘Them/Cat?’). Unrelated word pairs could either be preceded by the unspecific cue (50%) or either of the specific cues (25% each). After a 600 ms delay following cue presentation, prime and target word were presented in sequential order for 600 ms each, separated by a delay period of 600 ms. The nonius lines stayed on screen throughout cue, prime and target presentation, as well as the delay periods. At the end of each trial, nonius lines were presented for 900 ms, followed by a blank screen of 1200 ms. Trials were presented in pseudo-random order, such that transition probabilities between conditions were equal. Participants were instructed to respond as quickly and accurately as possible by pressing a key with their left or right index finger whether prime and target word were semantically related or unrelated.

**Figure 2.**
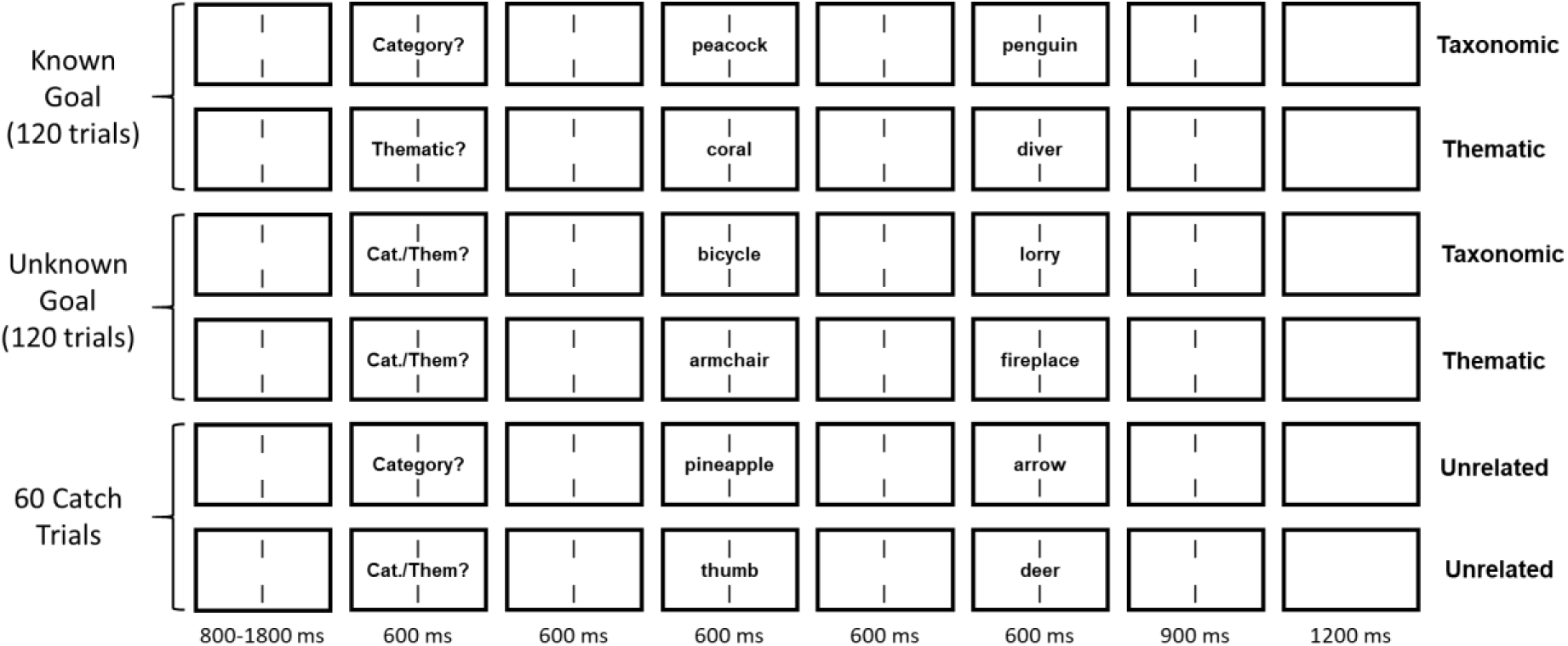
Experimental procedure. Each trial started with the presentation of nonius lines for a jittered interval between 800 and 1800 ms, followed by the presentation of a cue, a prime and a target stimulus for 600 ms each, separated by intervals of 600 ms. The nonius lines remained on screen the entire time, including a 900 ms interval following target offset, after which a blank screen was presented for 1200 ms. Prime and target could be taxonomically related, thematically related, or unrelated. Half of the taxonomically and thematically related trials were known goal trials, i.e., the cue indicated if they were taxonomically (‘Category?’) or thematically related (‘Thematic’). The other half were unknown goal trials, which were preceded by an unspecific cue (‘Them/Cat?’).

At the beginning of the study, participants were informed about the difference between taxonomic and thematic relations and performed a short training procedure of four trials in which they had to respond by button press if two simultaneously presented words shared a taxonomic or thematic relation and received feedback on every trial. Then, they completed a practice run of the main task including twelve trials with feedback on total accuracy presented at the end. If necessary, each training procedure was repeated until an accuracy of at least 75 % was reached. Trials from the training procedures were not used in the main experiment. After a subset of the behavioural data had been collected, we slightly changed the instruction at the start of the experiment: Participants were now explicitly informed that they would be presented with a higher number of related than unrelated word pairs. Of our included sample, nine participants were informed about this while eleven were not.

For the MEG study, stimuli were presented using Psychopy (version 3.1.5; Peirce et al., 2019) and projected onto a screen at a distance of ca. 90 cm of the participants. A neutral density filter was used to reduce glare. Presentation durations were identical to the behavioural study. Participants were instructed to respond as quickly and accurately as possible by pressing a key with their left or right index finger only when prime and target word were not semantically related. This approach prevented contamination of the MEG signal during related trials with the motor response. Response hands were counterbalanced across participants. Trials involving button presses were excluded from MEG data analysis. The MEG study was divided into four runs. Prior to the first run, participants completed a practice run of 25 trials with feedback on overall accuracy presented at the end. Training was repeated if the accuracy was less than 75 %. Participants were informed at the start of the experiment that there would be a higher number of related than unrelated word pairs. On the day before the MEG study, participants were informed about the difference between taxonomic and thematic relations via an online Zoom meeting, which was reiterated on the day of the MEG study.

### 2.4 Behavioural analysis

Statistical analyses of the behavioural study were performed using R, version 4.0.4, 2021-02-15 (R Development Core Team, 2008) and focused on the 240 semantically related trials selected for the MEG study. Log-transformed response times of correct responses were investigated in linear mixed models (LMMs) using the lmerTest package (Kuznetsova et al., 2017). For accuracies, we assumed a binary distribution for the outcome and therefore used generalized linear mixed models (GLMMs) with a logit link function. (G)LMMs included the fixed effects of task knowledge (known vs. unknown goal), semantic relation (taxonomic vs. thematic), and semantic similarity (based on word2vec), as well as two-way interactions of task knowledge with semantic relation and semantic similarity. The main models controlled for fixed effects of trial order and instruction (after the first 11 included participants, the instruction explicitly informed participants that there would be more related than unrelated word pairs), as well as random effects of participant, prime, and target. In addition, more complex models also controlled for fixed effects of the prime’s and target’s word length, word frequency, and imageability. Fixed effects were considered significant based on an absolute t-value above 2 in linear mixed models, and based on a p-value below 0.05 in generalized linear mixed models. For visualization of partial effects in the response time analysis, we used the remef package (Hohenstein & Kliegl, 2015).

### 2.5 MEG data acquisition

MEG data were acquired using a 248-sensor Magnes3600 system (4-D Neuroimaging, San Diego, California) at a sampling frequency of 2,000 Hz. Five sensors were disabled or excluded from analysis due to technical issues. External noise detected by the MEG reference channels was removed from the data. Five head localization coils recorded participants’ head positions relative to the sensor array at the beginning and end of each experimental run. Coil positions and participants’ head shapes were recorded with a 3D digitizer (Fastrak Polhemus) before MEG data acquisition to improve co-registration with the anatomical magnetic resonance (MR) images. For one participant, the recorded head shape was not saved due to technical issues. In this case, co-registration was based on landmarks (nasion, left and right pre-auricular points) identified from the anatomical MR image.

### 2.6 Structural MR image acquisition

Structural T1-weighted MR images were acquired with a 3 Tesla Siemens Prisma scanner (Siemens Medical Systems) using a 3D MPRAGE sequence (176 slices, 1×1×1 mm voxel size, 2.3 s TR, 2.26 s TE, 8° flip angle).

### 2.7 MEG preprocessing

The MEG data analysis focused on related word pairs. MEG data were preprocessed using FieldTrip (Version 06/2021; http://fieldtrip.fcdonders.nl; Oostenveld et al., 2011) under MATLAB (version 2019a, The MathWorks Inc., Natick, MA). MEG data were segmented into epochs containing a 700 ms baseline and the time windows of cue, prime, and target presentation, plus an additional 380 ms after target offset to allow for sufficient data for time-frequency analysis. Thus, epochs were lasting from −700 ms to 2,180 ms with respect to cue onset. 50 Hz line noise was removed using a discrete Fourier transform filter. Trials which included button presses at any time point from –700 ms until the onset of the next trial were excluded from analysis. In addition, trials which included sensor jump, muscle, or eye movement artifacts in the time period from −700 ms to 2,180 ms were excluded from analysis. Trials with artifacts in the baseline time window were excluded completely, while trials that only contained artifacts in the prime but not target presentation time window and vice versa were only excluded from analysis of the respective time window. Artifacts were identified using the standard FieldTrip routine for automatic artifact detection. Filters applied to the data prior to artifact detection were a 9th order median filter for jump artifacts, a 7th order Butterworth IIR filter (110-140 Hz) for muscle artifacts, and a 3rd order Butterworth IIR filter (2-15 Hz) for eye movement artifacts. Eye movement artifact detection was based on nine frontal sensors, while muscle and jump artifact detection took all sensors into account. The filtered data were *z*-transformed and averaged across sensors. Trials were rejected if, for any time point, the z-value exceeded a threshold of z = 40 for jump artifacts, z = 30 for muscle artifacts, and z = 10 for eye movement artifacts. A visual inspection confirmed this procedure was sufficient to remove artifacts. Out of 120 trials per condition, 85 trials on average per participant and condition (Known condition word 1: 85 trials, Known condition word 2: 85.7 trials, Unknown condition word 1: 84.8 trials, Unknown condition word 2: 84 trials) were retained for further analyses (range: 41 to 110).

### 2.8 MEG source localization

Source localization was performed separately for the first half (runs 1 and 2) and second half (runs 3 and 4) of the experiment, in order to minimize the effects of participants’ head movements over the course of the experiment. On average, the median difference in head position from the beginning of the first to the beginning of the third run or from the beginning of the third to the end of the fourth run across participants was 8 mm (range: 0.8 to 34.4 mm). Source localization was performed at the single-trial level using linearly constrained minimum variance (LCMV) spatial filters (Van Veen et al., 1997). The data covariance matrix was estimated based on all trials. Forward models were based on T1-weighted images co-registered and warped to an MNI-space template that comprised 1,619 voxels within the brain at 10 mm resolution. A single shell forward model of the inner surface of the skull (Nolte, 2003) was used for lead field computation. Spatial filters were computed based on the optimal dipole orientation that maximized spatial filter output, which was identified using singular value decomposition. Single-trial source power based on the precomputed spatial filters was then estimated for each voxel location.

### 2.9 Time-frequency analysis

Time-frequency analysis focused on total oscillatory power between 4 and 50 Hz, i.e., including theta (4-7.9 Hz), alpha (8-12.9 Hz), beta (13-29.9 Hz) and low gamma (30-50 Hz) ranges, which are sensitive to meaning-based contexts (Prystauka and Lewis, 2019), semantic task manipulations (Rahimi et al., 2022), and the strength of the association between words (Teige et al., 2018; 2019). A time-frequency transformation was performed on source power estimates using a Hanning taper with three cycles per time window, a frequency resolution of 1 Hz and a temporal resolution of 25 ms. A relative baseline correction using the time interval from –300 to –25 ms before the onset of the cue (i.e., –1,500 to –1,225 ms before the onset of the first word) was performed by dividing the oscillatory power during word presentation through the power during the baseline interval. The pre-cue interval was chosen as baseline in line with previous studies (e.g., McNab et al, 2007; Eisenhauer et al., 2022), as this is the only period without semantic retrieval and since separate baselines for both words may reduce anticipation effects for the second word which may already begin before its onset (e.g., Fruchter et al., 2015; Eisenhauer et al., 2022). For each participant and condition, time-frequency estimates were averaged across trials and the two halves of the experiment in three regions of interest (ROIs): Based on the fMRI study by Zhang et al. (2021), the left IFG cluster showing a task knowledge effect, as well as the bilateral visual cortex cluster, overlapping with the occipital poles and primary visual cortex, showing an interaction between task knowledge and word order were selected as ROIs. In addition, we selected the left pMTG cluster (which peaks in pMTG and posterior inferior temporal gyrus) implicated in semantic control from a recent fMRI meta-analysis (Jackson, 2021). These sites were selected to investigate the relationship between anterior and posterior nodes of the semantic control network (left IFG vs. pMTG) and the interaction of these regions with visual cortex, which passes on visual features of the words to higher order brain regions, in different phases of the paradigm, when the nature of the semantic relationship was known in advance or not. ROIs were defined based on masks provided by the original studies (Jackson, 2021; Zhang et al., 2021).

In each ROI, time-frequency power estimates in the 600 ms time window during word presentation were compared between conditions using cluster-based permutation tests (Maris and Oostenveld, 2007) for dependent samples, corrected for multiple comparisons across time points (0 to 600 ms with respect to word onset) and frequencies (4 to 50 Hz) at cluster level. One analysis included the 2×2 interaction between task knowledge (known vs. unknown) and word order (word 1 vs. 2), using the permANOVA function by Helbling (2015; https://github.com/sashel/permANOVA/) combined with Fieldtrip (version 20130106). A separate analysis, focused on the presentation time window of the second word, investigated the binary effects of high versus low semantic similarity (word2vec) based on a median split. For this analysis, the ten percent of trials closest to the median, identified separately for each knowledge condition and participant, were excluded from analysis. Clusters were defined as spectrally and temporally adjacent samples with F– or t-values exceeding an uncorrected α-level of 0.05 (0.025 at each tail in the case of t-values). The cluster-level statistic was calculated using the standard approach, i.e., taking the sum of F– or t-values within a cluster (Maris and Oostenveld, 2007). Empirical cluster-level statistics were compared to the distribution of cluster-level statistics obtained from Monte Carlo simulations with 1,000 permutations, in which condition labels were randomly exchanged within each participant. Original cluster-level statistics larger than the 95th percentile (below the 2.5^th^ and above the 97.5^th^ percentile in case of t-tests) of the distribution of cluster-level statistics obtained in the permutation procedure were considered significant.

### 2.10 Spectral Granger Causality

Source power estimates were baseline corrected by subtracting the power from the interval between –300 to 0 ms before the onset of the cue. A fast Fourier transform between 0 and 100 Hz was performed on source power estimates using multitapers with a frequency resolution of 3.3 Hz for three overlapping time windows of interest: 0 to 300 ms, 150 to 450 ms, and 450 to 600 ms with respect to word onset (see Schaum et al., 2021, for a similar approach). Multivariate conditional spectral Granger Causality (Granger, 1969; Geweke, 1982) was used to assess the direction of information flow between ROIs. This method reveals a directed connection from brain region X to brain region Y if the prediction of the present time series of Y based on its past time series can be improved by adding the past time series of X (Dhamala et al., 2008), conditional on the time series in brain region Z. This method allowed us to infer the directed connectivity between pairwise combinations of ROIs, while taking into account potential connections with the third ROI. The spectral Granger Causality analysis was performed on the single-trial Fourier transformed data combined across the two halves of the experiment for each pairwise combination of the three ROIs, obtaining feedforward and feedback spectral connectivity estimates per participant, condition, and time window.

For each combination of a pair of ROIs, a connectivity direction (feedforward or feedback), and a time window, the 2×2 interaction between task knowledge (known vs. unknown) and word order (word 1 vs. 2) was assessed using a cluster-based permutation test (Maris and Oostenveld, 2007) and the permANOVA function by Helbling (2015; https://github.com/sashel/permANOVA/) combined with Fieldtrip (version 20130106). In a separate analysis, we investigated the effect of semantic similarity (based on word2vec) on spectral Granger Causality estimates during the presentation of the second word, using the median-split approach as described for the time-frequency analysis. Multiple comparison correction across frequencies (4 to 50 Hz) was performed at the cluster level. Resulting p-values were then Bonferroni-corrected for the number of tests across ROI combinations, connectivity directions, and time windows (18 tests in total). Clusters were defined as spectrally adjacent samples with F– or t-values exceeding an uncorrected α-level of 0.05 (0.025 at each tail in the case of t-values). The cluster-level statistic was calculated by taking the sum of F-values within a cluster (Maris and Oostenveld, 2007). Empirical cluster-level statistics were compared to the distribution of cluster-level statistics obtained from Monte Carlo simulations with 5,000 permutations, in which condition labels were randomly exchanged within each participant. Original cluster-level statistics larger than the 95th percentile (below the 2.5^th^ and above the 97.5^th^ percentile in case of t-tests) of the distribution of cluster-level statistics obtained in the permutation procedure were considered significant.

### 2.11 Code and Data Accessibility

All analysis code is publicly available at the Open Science Framework (https://osf.io/j9pz4/). The conditions of our ethics approval do not permit public archiving of the raw MEG and MRI data supporting this study. Readers seeking access to this data should contact the lead author, Susanne Eisenhauer, the PI Professor Beth Jefferies, or the local ethics committee at the Department of Psychology and York Neuroimaging Centre, University of York. Access will be granted to named individuals in accordance with ethical procedures governing the reuse of sensitive data. Specifically, the following conditions must be met to obtain access to the data: approval by the Department of Psychology and York Neuroimaging Research Ethics Committees and a suitable legal basis for the release of the data under GDPR.

## 3 Results

### 3.1 Behaviour

In the behavioural study, we found significant main effects of task knowledge and semantic similarity, reflecting faster responses when the semantic goal was known vs. unknown (Fig. 3a), as well as when the two words were more strongly linked (Fig. 3b; Table 2). The main effect of trial order also reached significance, indicating that response times increased throughout the experiment. There was no main effect of semantic relation (taxonomic vs. thematic) and no interactions between task knowledge and either semantic relation or semantic similarity. The same pattern of results was found in a more complex linear model (based on the subset of trials for which imageability ratings were available) that also controlled for the effects of prime and target word length, word frequency, and imageability (Table 3).

**Figure 3.**
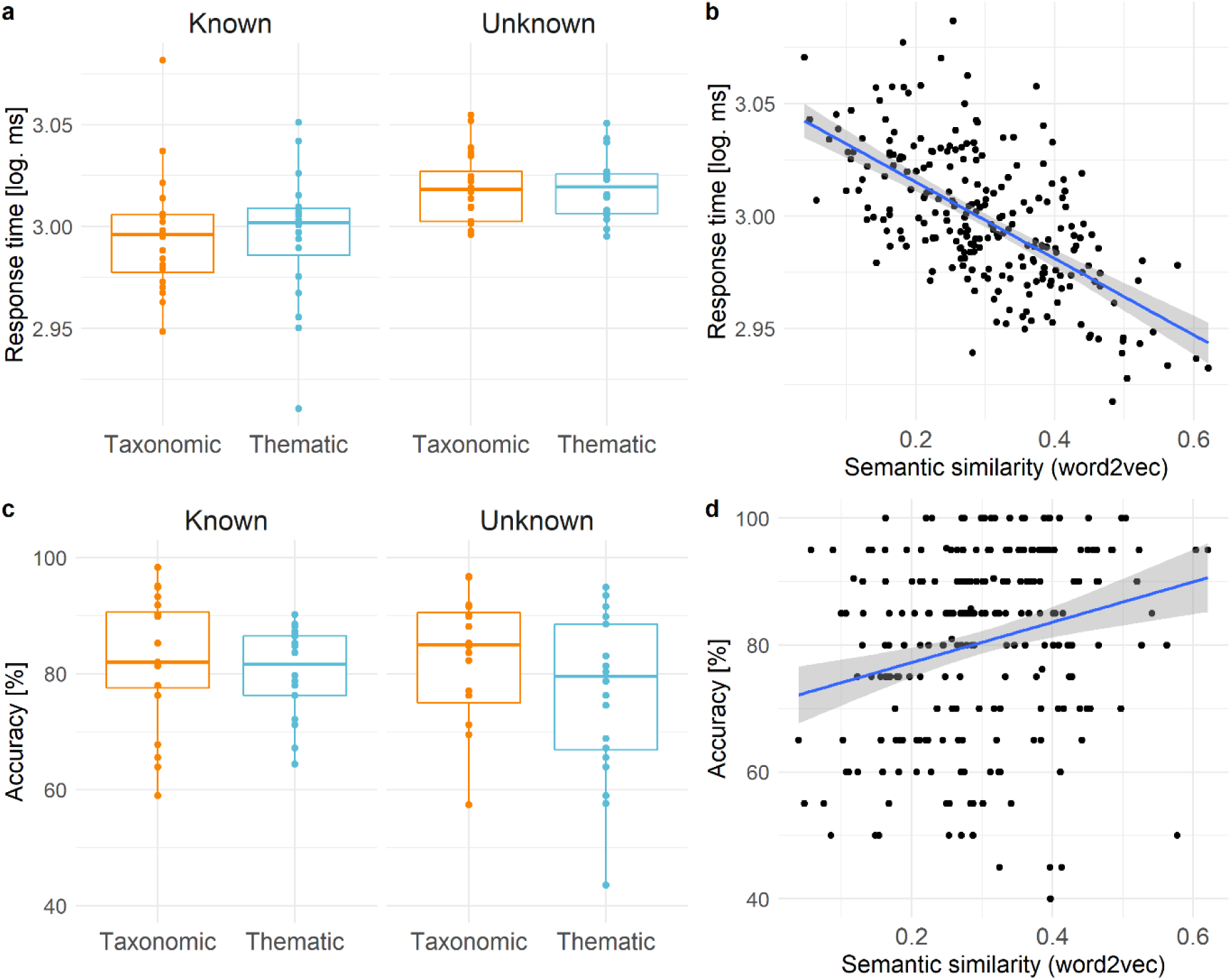
Response times and accuracy for each condition. (a, c) as well effects of semantic similarity based on word2vec (b, d). In a and b, partial effects from linear mixed models are shown. Each dot represents a word averaged across participants.

**Table 2.** Results of the simple linear mixed model of behavioural response times.

|  | FE | SE | *t* |
| --- | --- | --- | --- |
| Intercept | **2.995** | **0.024** | **126.65** |
| Task Knowledge | **0.023** | **0.006** | **4.05** |
| Semantic Relation | 0.003 | 0.007 | 0.43 |
| Semantic Similarity (word2vec) | **-0.020** | **0.004** | **-5.54** |
| Trial Order | **-0.024** | **0.002** | **-11.20** |
| Instruction | -0.015 | 0.023 | -0.65 |
| Task Knowledge x Semantic Relation | -0.010 | 0.008 | -1.22 |
| Task Knowledge x Semantic Similarity (word2vec) | 0.003 | 0.004 | 0.84 |
| Bold numerals indicate significant effects, i.e., $\vert t\vert > 2$. FE = fixed effect estimates; SE = standard error. | | | |

**Table 3.**
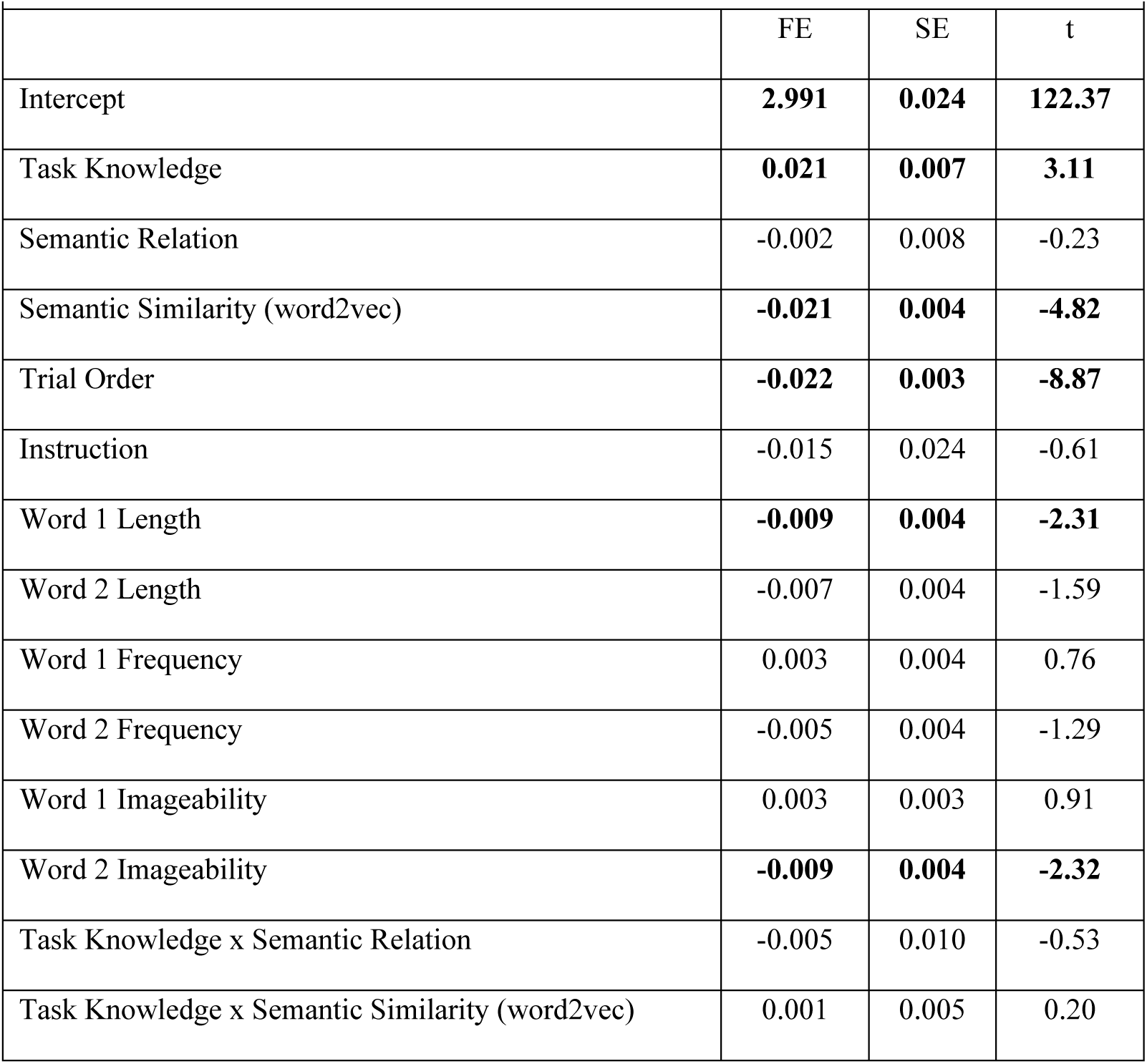
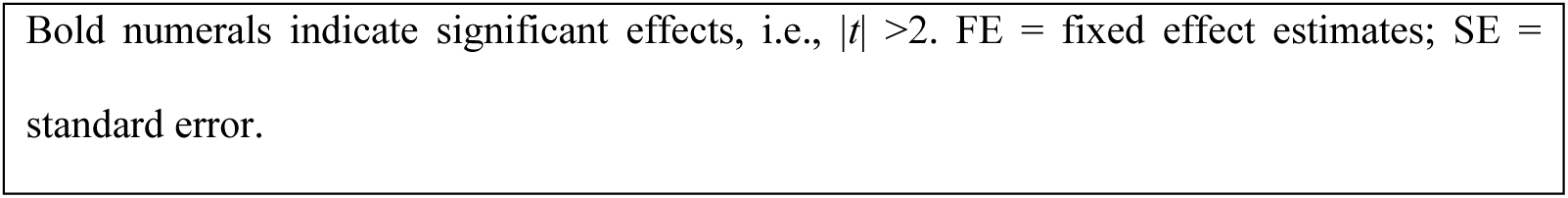
Results of the complex linear mixed model of behavioural response times.

|  | FE | SE | t |
| --- | --- | --- | --- |
| Intercept | **2.991** | **0.024** | **122.37** |
| Task Knowledge | **0.021** | **0.007** | **3.11** |
| Semantic Relation | -0.002 | 0.008 | -0.23 |
| Semantic Similarity (word2vec) | **-0.021** | **0.004** | **-4.82** |
| Trial Order | **-0.022** | **0.003** | **-8.87** |
| Instruction | -0.015 | 0.024 | -0.61 |
| Word 1 Length | **-0.009** | **0.004** | **-2.31** |
| Word 2 Length | -0.007 | 0.004 | -1.59 |
| Word 1 Frequency | 0.003 | 0.004 | 0.76 |
| Word 2 Frequency | -0.005 | 0.004 | -1.29 |
| Word 1 Imageability | 0.003 | 0.003 | 0.91 |
| Word 2 Imageability | **-0.009** | **0.004** | **-2.32** |
| Task Knowledge x Semantic Relation | -0.005 | 0.010 | -0.53 |
| Task Knowledge x Semantic Similarity (word2vec) | 0.001 | 0.005 | 0.20 |

Bold numerals indicate significant effects, i.e., $|t| > 2$ . FE = fixed effect estimates; SE = standard error.

Concerning response accuracy, the simple general linear model revealed a significant interaction between task knowledge and semantic relation (Table 4). Accuracy was higher for known vs. unknown trials in thematic but not taxonomic word pairs (Fig. 3c, Table 5). Moreover, accuracy was significantly higher for word pairs of increasing semantic similarity, irrespective of task knowledge (Fig. 3d). In the more complex linear model controlling for prime and target word characteristics (based on the subset of trials for which imageability ratings were available), a similar main effect of semantic similarity was found; however, the interaction between task knowledge and semantic relation, as well as the main effect of task knowledge, did not reach significance (Table 6). To sum up, the behavioural findings indicated that semantic relatedness decisions were facilitated both when semantic goals were known in advance, as well as when word pairs were more semantically similar.

**Table 4.** Results of the simple generalized linear mixed model of behavioural accuracies.

| | FE | SE | $z$ | $p$ |
| --- | --- | --- | --- | --- |
| Intercept | **1.758** | **0.174** | **10.09** | **< 2e-16** |
| Task Knowledge | 0.041 | 0.113 | 0.36 | 0.7162 |
| Semantic Relation | 0.016 | 0.155 | 0.10 | 0.9178 |
| Semantic Similarity (word2vec) | **0.286** | **0.080** | **3.60** | **0.0003** |
| Trial Order | 0.037 | 0.040 | 0.92 | 0.3580 |
| Instruction | 0.076 | 0.138 | 0.55 | 0.5832 |
| Task Knowledge x Semantic Relation | **-0.342** | **0.159** | **-2.15** | **0.0312** |
| Task Knowledge x Semantic Similarity (word2vec) | -0.077 | 0.081 | -0.96 | 0.3392 |

Bold numerals indicate significant effects, i.e., $p < 0.05$ . FE = fixed effect estimates; SE = standard error.

**Table 5.** Post-hoc results of the simple generalized linear mixed model of behavioural accuracies for trials with thematic and trials with taxonomic relations.

|  | FE | SE | *z* | *p* |
| --- | --- | --- | --- | --- |
| **Thematic relation** |  |  |  |  |
| Intercept | **1.675** | **0.164** | **10.25** | **<2e-16** |
| Task Knowledge | **-0.255** | **0.107** | **-2.39** | **0.017** |
| Semantic Similarity (word2vec) | 0.207 | 0.108 | 1.92 | 0.056 |
| Trial Order | -0.008 | 0.054 | -0.16 | 0.876 |
| Instruction | 0.101 | 0.132 | 0.77 | 0.442 |
| Task Knowledge x Semantic Similarity (word2vec) | 0.139 | 0.109 | 1.27 | 0.204 |
| **Taxonomic relation** |  |  |  |  |
| Intercept | **1.955** | **0.214** | **9.12** | **< 2e-16** |
| Task Knowledge | -0.007 | 0.116 | -0.06 | 0.954 |
| Semantic Similarity (word2vec) | **0.381** | **0.116** | **3.28** | **0.001** |
| Trial Order | 0.092 | 0.060 | 1.53 | 0.126 |
| Instruction | 0.018 | 0.186 | 0.10 | 0.922 |
| Task Knowledge x Semantic Similarity (word2vec) | **-0.349** | **0.119** | **-2.95** | **0.003** |
| Bold numerals indicate significant effects, i.e., $p < 0.05$. FE = fixed effect estimates; SE = standard error.<br>For thematic trials, the full model did not converge, and results are reported for a model without the random effect of the prime, which explained the least variance in the data. |  |  |  |  |

**Table 6.** Results of the complex generalized linear mixed model of behavioural accuracies.

|  |  |  |  |  |
| --- | --- | --- | --- | --- |
|  | FE | SE | *z* | *p* |
| Intercept | 1.803 | 0.184 | 9.82 | < 2e-16 |
| Task Knowledge | -0.043 | 0.134 | -0.32 | 0.748 |
| Semantic Relation | 0.006 | 0.180 | 0.04 | 0.971 |
| Semantic Similarity (word2vec) | **0.377** | **0.097** | **3.89** | **0.0001** |
| Trial Order | -0.029 | 0.049 | -0.60 | 0.552 |
| Instruction | 0.101 | 0.139 | 0.73 | 0.467 |
| Word 1 Length | -0.015 | 0.084 | -0.17 | 0.863 |
| Word 2 Length | 0.137 | 0.096 | 1.43 | 0.154 |
| Word 1 Frequency | 0.000 | 0.088 | 0.00 | 0.998 |
| Word 2 Frequency | 0.079 | 0.086 | 0.92 | 0.359 |
| Word 1 Imageability | 0.140 | 0.073 | 1.91 | 0.056 |
| Word 2 Imageability | **0.222** | **0.084** | **2.64** | **0.008** |
| Task Knowledge x Semantic Relation | -0.198 | 0.197 | -1.00 | 0.315 |
| Task Knowledge x Semantic Similarity (word2vec) | -0.069 | 0.101 | -0.68 | 0.495 |
| Bold numerals indicate significant effects, i.e., $p < 0.05$. FE = fixed effect estimates; SE = standard error. | | | | |

In the MEG study, participants performed the task with high accuracy, correctly responding to 93.7% of unrelated trials, and correctly refraining from a response to 93.8% of known and 93.6% of unknown goal trials.

### 3.2 Time-frequency representations

We investigated the interaction of task knowledge and word order, using cluster-based permutation tests in three ROIs: the left IFG, left pMTG, and bilateral visual cortex (Fig. S1). The presentation of the two words elicited synchronized oscillatory responses relative to the pre-cue baseline in all three ROIs (Fig. S1). Whole-brain oscillatory power for the effect of task knowledge is shown in Figure S2. Main effects of word order were found in all three ROIs and are shown in Figure S3. Main effects of task knowledge and its interactions with word order are described below.

In left IFG, a main effect of task knowledge was found in two clusters (Fig. 4, top). In the theta/alpha band, relative power was higher in the known than in the unknown goal condition from ∼100 to 350 ms after word onset. In the gamma band at 225 ms, the opposite pattern with higher power in the unknown than in the known goal condition was found.

**Figure 4.**
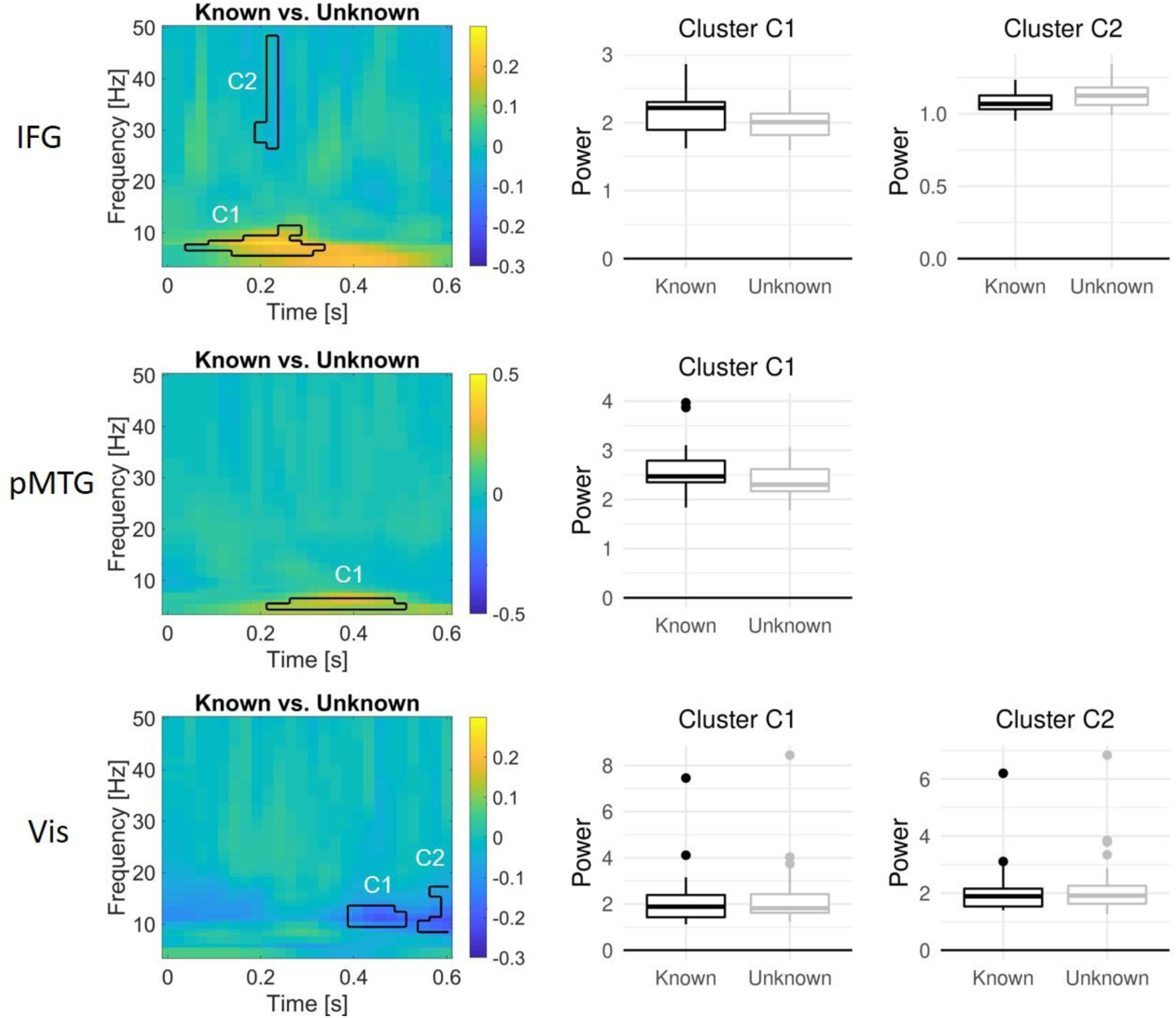
Oscillatory dynamics of the main effect of task knowledge in left IFG (top), left pMTG (middle), and bilateral visual cortex (bottom), locked to word onset. Time-frequency plots show the difference in power (relative to the pre-cue baseline) between known and unknown trials, averaged across the first and second word, in significant clusters. Boxplots show the oscillatory power in each significant cluster for known and unknown trials across participants. Raw oscillatory responses to the first and second word are shown in Figure S1. Whole-brain oscillatory responses for the difference between known and unknown trials are shown in Figure S2. Main effects of word order are shown in Figure S3. A follow-up analysis comparing the strength of the known > unknown effect in IFG and pMTG across time revealed main effects of knowledge but no interaction with ROI (Fig. S4).

In left pMTG, a main effect of task knowledge was found from ∼200 to 500 ms, indicating higher power in the known than in the unknown goal condition in the theta band (Fig. 4, middle). Thus, this effect occurred in a similar frequency range, and seemingly later but partly overlapping time window, as the known over unknown effect in the IFG. Since the exact timing of cluster-based permutation statistics cannot be inferred from the temporal extent of the cluster (Sassenhagen and Draschkow, 2019), we ran a follow-up analysis assessing the interaction between the knowledge effect and ROI (IFG vs. pMTG) at each time point to reveal potential points in time when the knowledge effect is stronger in one region over the other. However, this analysis only revealed main effects of knowledge and no interaction (Fig. S4), and thus did not provide evidence for a temporal separation of the effects in IFG and pMTG.

In bilateral visual cortex, main effects of task knowledge were found in two temporally adjacent clusters in the alpha/low beta range between 400 and 600 ms, indicating higher power in the unknown compared to the known goal condition (Fig 4, bottom). Furthermore, bilateral visual cortex showed interactions between task knowledge and word order in three clusters (Fig. 5). Within the first 200 ms, alpha/low beta power increased from the first to the second word, particularly within the unknown condition. In two clusters from ∼450 to 600 ms, high alpha to low gamma power decreased from the first to the second word, and remained higher in the unknown than in the known condition. These two later clusters highly overlapped with the two clusters showing a main effect of task knowledge.

**Figure 5.**
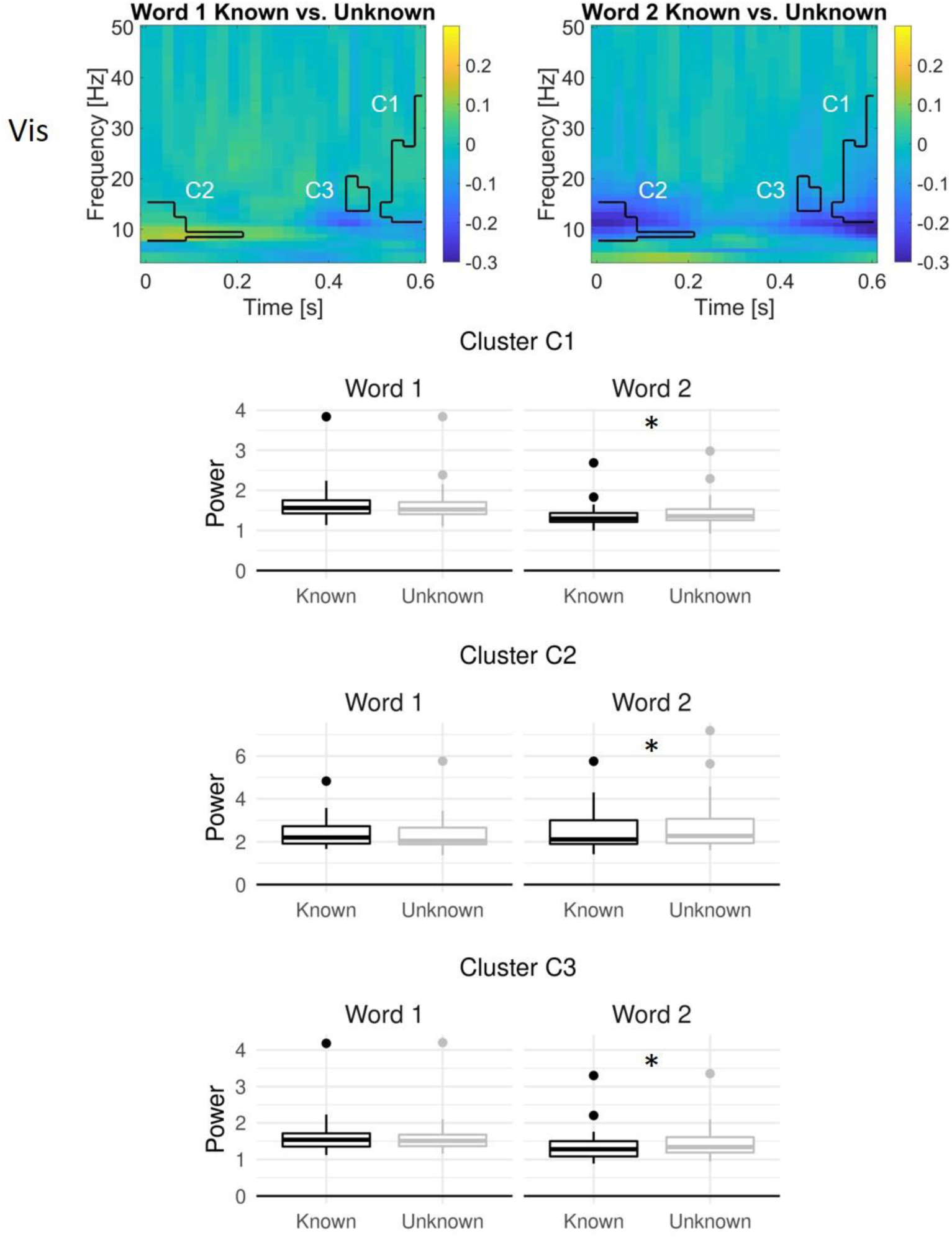
Oscillatory dynamics of the interaction between task knowledge and word order in bilateral visual cortex, locked to word onset. Time-frequency plots show the difference in power (relative to the pre-cue baseline) between known and unknown trials for word 1 (left) and word 2 (right) in significant clusters. Boxplots show the oscillatory power in each significant cluster for known and unknown trials in word 1 and word 2 across participants. Asterisks indicate post-hoc significant differences between the known and unknown goal conditions at the second word only. No significant interactions were found for the other ROIs.

Furthermore, we investigated effects of the semantic similarity between the two related words in a trial, based on word2vec. Significant effects of semantic similarity were found in pMTG from 350 to 550 ms (peaking at 425 ms, 12 Hz; Fig. 6, top), as well as in visual cortex from 0 to ∼550 ms (peaking at 50 ms, 10 Hz; Fig. 6, bottom). In both regions, alpha/low beta power was higher for word pairs with lower semantic similarity.

**Figure 6.**
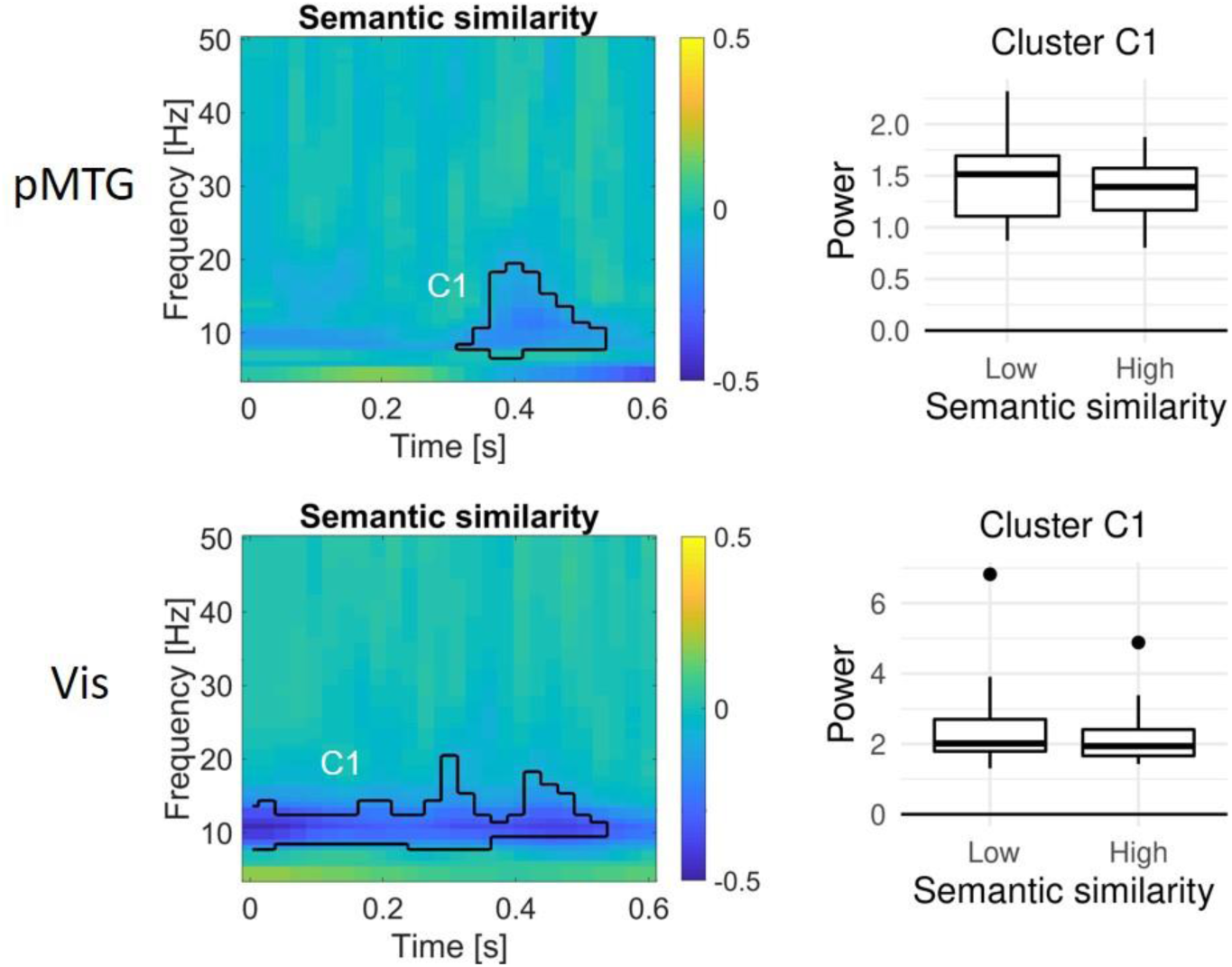
Oscillatory dynamics of the effect of semantic similarity (word2vec) during the processing of the second word in left pMTG, and bilateral visual cortex, locked to word onset. Time-frequency plots show the difference in power (relative to the pre-cue baseline) between word pairs of high vs. low semantic similarity in significant clusters. Boxplots show the oscillatory power in each significant cluster for word pairs of high vs. low semantic similarity across participants. No significant effect of semantic similarity was found in the left IFG.

To summarize our findings (Fig. 7), the earliest observed knowledge effects were increases in theta/alpha band activity in left IFG, peaking at 7 Hz and 200 ms after word onset (across both words). At a similar time (∼225 ms), IFG high beta/gamma activity (peaking at 33 Hz) showed the reverse pattern with higher power in the unknown goal condition. Around 350 ms post word onset, theta band activity in left pMTG (peaking at 6 Hz) resembled the pattern observed earlier in IFG, i.e., higher power in the known condition. Bilateral visual cortex only demonstrated significant knowledge effects for the second word. Here, greater power for the unknown goal condition was identified around stimulus onset, in alpha/beta frequencies (peaking at 10 Hz), and around 600 ms, within the beta/gamma range (peaking at 31 Hz). When participants made decisions about less similar word pairs, there was higher alpha/beta power in bilateral visual cortex extending throughout much of the trial after presentation of the second word. A similar pattern was found in the left pMTG, peaking at around 425 ms.

**Figure 7.**
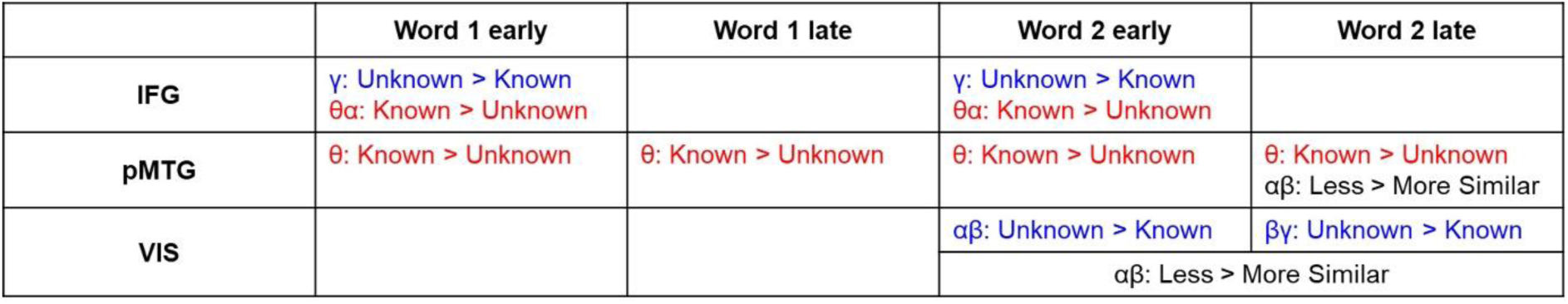
Summary of oscillatory dynamics. For each ROI, differences in oscillatory power between known and unknown goal trials in early (from 0 to 300 ms) and late time windows (from 300 to 600 ms) during the presentation of words 1 and 2 are presented. In case of significant main effects of knowledge in the IFG and pMTG, follow-up paired t-tests on oscillatory power averaged within the significant cluster confirmed that knowledge effects were significant (p < 0.05) for both word 1 and word 2 in most cases, except for the theta/alpha effect in the IFG at word 2, which was marginally significant (p = 0.055). For word 2, differences between less and more semantically similar word pairs are presented additionally. The frequency band of observed effects is indicated. Blue: Unknown > Known. Red: Known > Unknown. Black: Less > More Similar.

### 3.3 Spectral Granger Causality

We used spectral Granger Causality to investigate task knowledge effects (Fig. 8) and their interaction with word order (Fig. 9) on the effective connectivity between our three ROIs in early, intermediate and late time windows after word onset. In the early time window from 0 to 300 ms, a significant interaction between task knowledge and word order was found for the connection from visual cortex to pMTG in the gamma band (Fig. 9b): While this connection significantly increased from word 1 to word 2 both for known and unknown trials, the effect was numerically stronger in the unknown condition. Additionally, a significant main effect of knowledge indicated that the connection from visual cortex to pMTG was stronger for known trials in the beta band (Fig. 8b). Furthermore, connectivity from left IFG to pMTG in the gamma band was stronger in the unknown than in the known goal condition at the second word only, indicated by a significant interaction between goal condition and word order (Fig. 9a).

**Figure 8.**
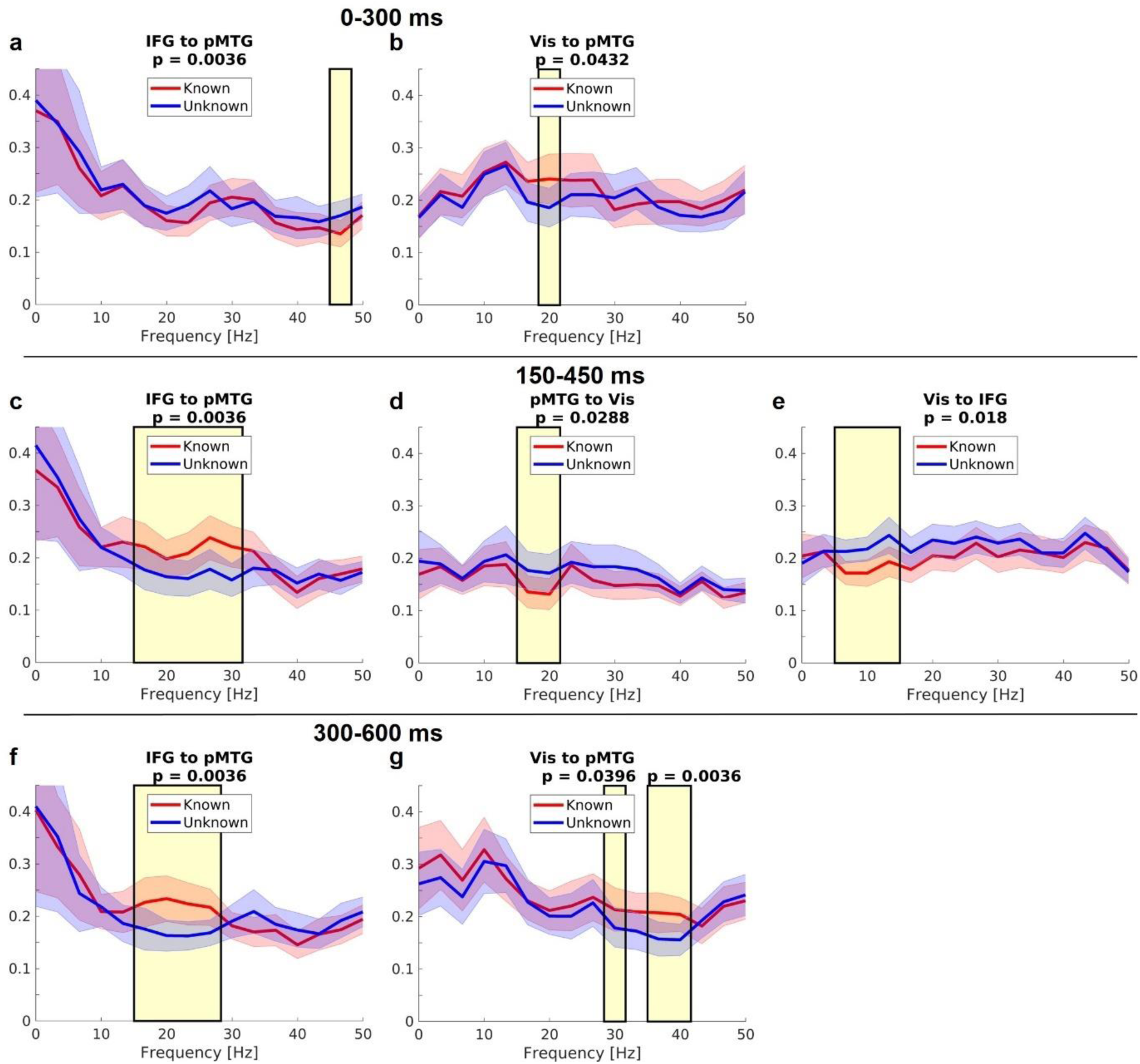
Main effects of task knowledge in spectral Granger causality estimates (averaged across word 1 and 2) in the time windows 0 to 300 ms, 150 to 450 ms, and 300 to 600 ms after word onset. Shading around the lines indicates standard errors. Yellow shading indicates significant frequency ranges. Corresponding Bonferroni-corrected p-values are shown. Only significant effects of task knowledge on connectivity are plotted.

**Figure 9.**
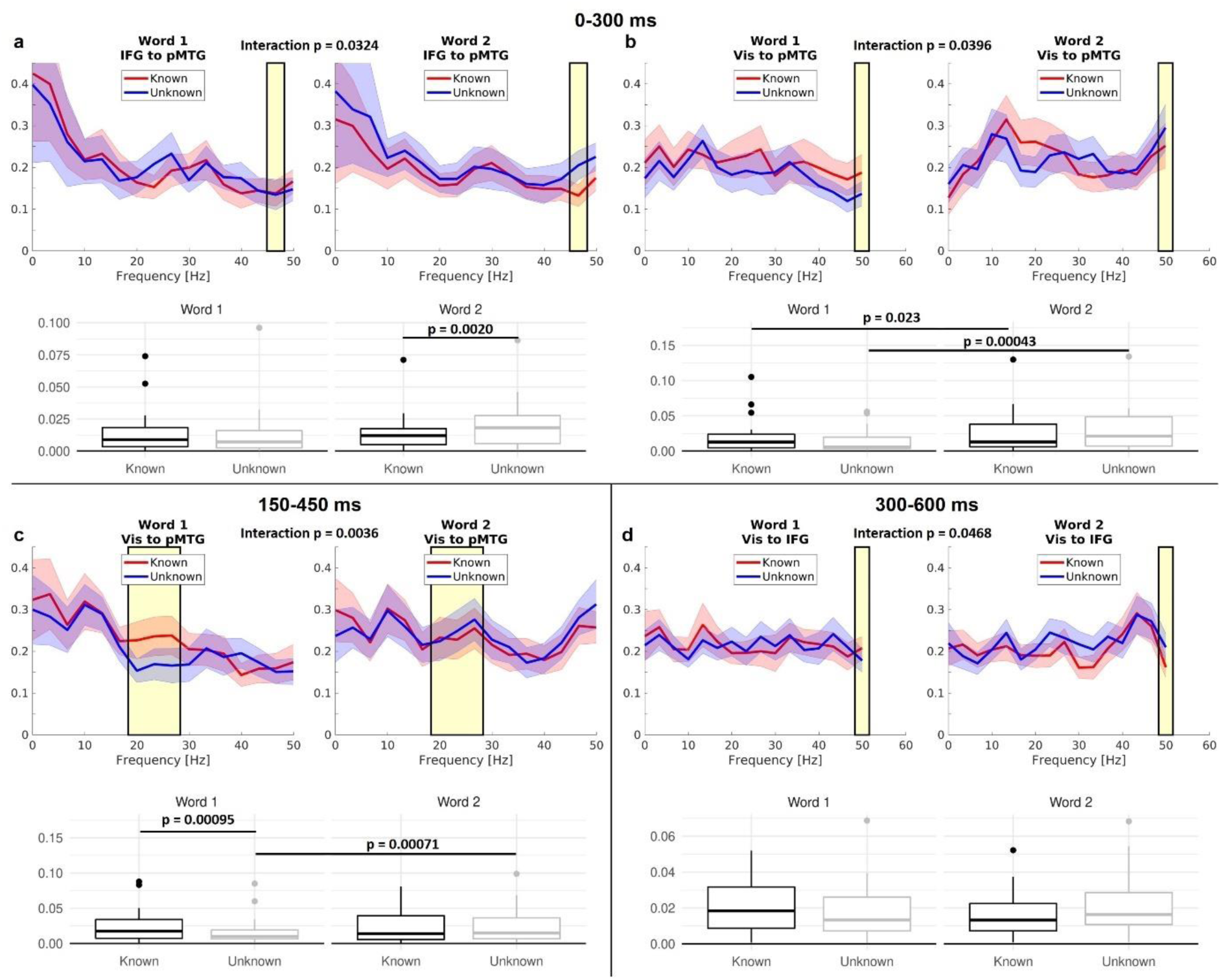
Interactions between task knowledge and word order in spectral Granger causality estimates in the time windows 0 to 300 ms, 150 to 450 ms, and 300 to 600 ms after word onset. Left, word 1; right, word 2. Shading around the lines indicates standard errors. Yellow shading in line plots indicates the significant frequency range. The corresponding Bonferroni-corrected p-values of the interactions are shown. Boxplots show the average power in the significant cluster across participants for each word and knowledge condition as well as p-values for significant contrasts from post-hoc tests. Only significant connectivity interactions are plotted.

In the intermediate time window from 150 to 450 ms, an interaction for the feedforward connection from visual cortex to pMTG emerged in the beta band (Fig. 9c), with stronger connectivity for known vs. unknown goals at the second word only, and stronger connectivity for the second vs. first word in the unknown condition. Feedback connectivity from IFG to pMTG in the beta band was stronger for known than unknown trials (Fig. 8c). Stronger connections in the unknown condition were found from pMTG to visual cortex in the beta band (Fig. 8d), and from visual cortex to IFG in the alpha band (Fig. 8e).

In the final time window from 300 to 600 ms, the connections from the IFG to pMTG in the beta band (Fig. 8f), as well as from visual cortex to pMTG in the high beta/gamma band (Fig. 8g), were stronger in the known than in the unknown goal condition. In addition, a significant interaction for the connection from visual cortex to the IFG emerged in the alpha/low beta band, however, post-hoc tests did not reveal any significant effects resolving this interaction (Fig. 9d). We did not observe significant effects of semantic similarity on Granger causality estimates in any of the investigated ROI pairs and time windows.

To summarize our key findings (Fig. 10), IFG provided top-down feedback to pMTG, which was stronger when retrieval goals were known in intermediate and late time windows, irrespective of word order, and stronger when goals were absent at the second word only, in the early time window. pMTG provided top-down feedback to visual cortex when goals were absent, irrespective of word order in the intermediate time window. Feedforward signalling from visual cortex to pMTG was stronger for known than unknown trials, in particular for the first word (in the intermediate time window).

**Figure 10.**
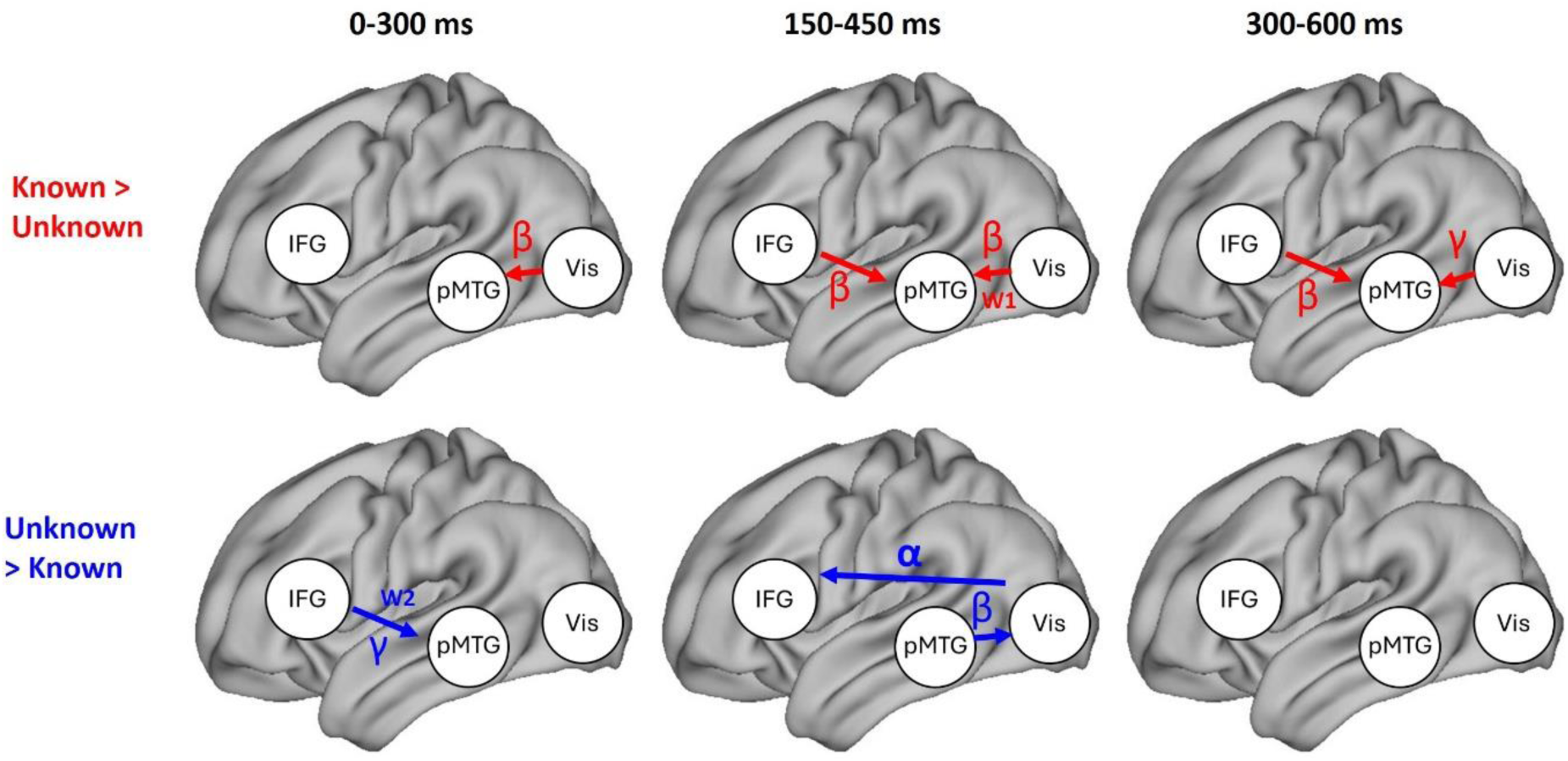
Effective connectivity based on spectral Granger causality estimates between left IFG, left pMTG and bilateral visual cortex in the time windows 0 to 300 ms, 150 to 450 ms, and 300 to 600 ms after word onset. Red arrows (top) show connections that are stronger in the known goal condition, while blue arrows (bottom) show connections that are stronger in the unknown goal condition. W1 and W2 indicate that the known vs. unknown difference was found at the first or the second word, respectively, based on a significant interaction between task knowledge and word order and follow-up comparisons. The frequency range in which significant effects were found is also indicated. Only connections with a significant main effect of task knowledge, or a significant effect of task knowledge at the first or second word following a significant interaction between task knowledge and word order, are shown.

## 4 Discussion

Faced with the abundant semantic information associated with each concept, we rely on semantic control mechanisms within the left IFG and pMTG for the efficient retrieval of features that are relevant to our current context and goals. The present investigation disentangles the roles of IFG and pMTG for controlled semantic cognition, indexed by their spectro-temporal profile and interactions with visual activity.

Both knowledge of the type of semantic relation relevant in a given trial, and stronger semantic similarity between words, enabled faster behavioural response times, showing that contextual information facilitates semantic cognition (e.g., Mandera et al., 2017; Eisenhauer et al., 2019; 2022). However, these factors differently affected MEG-measured oscillatory dynamics in two semantic control regions (IFG and pMTG), as well as in visual cortex.

Semantic similarity effects, i.e., higher oscillatory power for less similar words, were found in the alpha/beta band in visual cortex from word onset, and in the theta band in left pMTG peaking around 350 ms, the latter replicating previous findings (Teige et al., 2019). While both visual cortex and pMTG were sensitive to input-driven demands of semantic cognition, no significant differences were found in effective connectivity between more and less semantically similar word pairs.

Top-down guided semantic cognition, reflected in task knowledge effects, interacted with the bottom-up input, i.e., the order of the words, in visual cortex. This interaction was observed both in oscillatory power and in connectivity from visual cortex, with different underlying patterns: For trials with known versus unknown retrieval goals and during the whole time window of word presentation, visual cortex showed a sustained feedforward connection to left pMTG, which occurred in the beta band in early and intermediate time windows (0 to 450 ms), and in the high beta/gamma band in the later time window (300 to 600 ms). During the intermediate time window, this effect was present at the first word only, suggesting that knowledge of semantic retrieval goals allows for enhanced information transfer from visual to posterior temporal cortex already at the first word. However, alpha/beta power in visual cortex was reduced only at the second word when semantic goals were known in advance. This effect was found immediately after word onset, as well as towards the end of word presentation, suggesting that semantic goals facilitate visual processing both during the earliest stages of retrieval, as well as close in time to the semantic decision.

The observed interactions – with stronger connections from visual cortex to pMTG at the first word for known goals, but reduced power in visual cortex at the second word – resemble previous fMRI findings, showing enhanced activation for known goals at the first word, and for unknown goals at the second word (Zhang et al., 2021; Fig. 1a, right). The pattern in the present study suggests that known goals enable more rapid feature retrieval from the first word in visual cortex, which ultimately facilitates visual processing of the second word.

In semantic control regions, oscillatory dynamics for known versus unknown goals, irrespective of word order, were characterized by stronger theta/alpha power in left IFG and pMTG in partly overlapping time windows (peaking at 200 ms and ∼350 ms, respectively), as well as stronger beta/low gamma connectivity from IFG to pMTG in the intermediate and late time windows (150 to 600 ms). These findings indicate that the semantic control network does not only engage more when semantic cognition is demanding (i.e., when semantic associations are weaker, e.g., Snyder et al., 2011; Krieger-Redwood et al., 2015; 2023; Teige et al., 2019) but also when semantic goals are explicitly known and facilitate semantic decisions (Grindrod et al, 2008; Zhang et al., 2021; cf. Wang et al., 2021; Fig. 1a, left). Moreover, while previous studies found functional connections between IFG and pMTG at rest (Hallam et al., 2016; Wawrzyniak et al., 2017; Zhang et al., 2021) and during semantic cognition (Yue et al., 2013; Davey et al., 2016; Jackson et al., 2016), our findings reveal the directionality of this connection and its modulation by task knowledge.

We provide strong evidence for differential roles of anterior and posterior regions within the semantic control network. A recent computational model suggested that context-dependent semantic representations are activated through an intermediate network layer that receives constraints from a task-dependent control system and mutually interacts with the input system (Giallanza et al., 2024). Our observations are consistent with a view in which the IFG as control region constrains processing in pMTG (Lau et al., 2008), while pMTG integrates this top-down information with the bottom-up input (Fig. 1c, right). While we cannot disentangle the exact nature of the representation reflected by the knowledge effect in each region, the IFG may sustain both abstract task goals and goal-relevant semantic representations (Wang et al., 2021), and both might contribute to the observed pattern. Given the pMTG’s implication in the storage of semantic representations (Schwartz et al., 2011, Teige et al., 2019) and lexical-semantic access (Dikker et al., 2020), the integration of top-down and bottom-up information within pMTG potentially allows for the rapid activation of context-dependent semantic features.

In contrast, when semantic retrieval goals were unknown, irrespective of word order, IFG received direct input from visual cortex, which simultaneously received input from pMTG. While this pattern of information flow from pMTG to visual cortex to IFG was not predicted, it is consistent with a role for pMTG in constraining visual activation when absent contextual cues result in higher processing demands, in line with the role of this region in semantic control (Davey et al., 2016; Fig. 1c, left). Moreover, while IFG showed higher gamma power for both words around 200 ms, this region projected to pMTG only at the second word, i.e., after having received bottom-up information from visual cortex for the first word, in line with delayed activation of semantic representations when task goals were absent.

While knowledge effects on oscillatory power in semantic control regions preceded effects in visual cortex, which only occurred at the second word (in line with our hypothesis, Fig. 1a), connectivity findings revealed feedback from pMTG within the semantic control network to visual cortex only in the unknown condition. Thus, in contrast to our expectation (Fig. 1b), our findings do not provide evidence for direct gating of visual activation by semantic control regions when semantic retrieval goals are known. While a computational model postulated direct connections from control to input regions in task-dependent semantic cognition (Jackson et al., 2021), task knowledge in this model represented modality, e.g., auditory vs. visual. In contrast, knowing the type of semantic relation as in the present study may involve less direct feedback from control to input regions.

A limitation of the present study is the use of a simple semantic context with a relatively small variance in semantic similarity. Although this allowed good experimental control of semantic goals and reduced variability from the linguistic input, it remains to be seen if the same mechanisms generalize to more naturalistic contexts (Deniz et al., 2023). Since IFG couples with temporal cortex during object (Xu et al., 2020) and face recognition (Zhen et al., 2013) and reflects goals across various domains, including action perception (Wurm et al., 2014; Hrkać et al., 2015) and emotion regulation (Morawetz et al., 2016), our findings generate broad predictions about how IFG might modulate temporal activation in goal-dependent cognition across domains, which should be investigated in future research. Furthermore, the present findings cannot dissociate if reduced activation in visual cortex reflected facilitated recognition of letters/orthography (Eisenhauer et al., 2019), and/or facilitated retrieval of visual-semantic features of the corresponding object (Chiou et al., 2018; Zhang et al., 2021). Moreover, while previous studies varying linguistic context mainly found desynchronized alpha/beta oscillations (Prystauka and Lewis, 2019), we observed synchronization across all frequency bands. Possibly, the temporal onset predictability of the presented words based on the preceding cue may have driven synchronized responses.

The significant frequencies did not always line up between oscillatory power and connectivity. In the known condition, both IFG and pMTG showed higher theta power, while connectivity from IFG to pMTG was stronger in the beta band. Theta band effects in connectivity may be absent either because the power effects were overlapping in time, indicating that no strong directionality may underly these effects, or due to lower sensitivity of Granger causality to detect effects at lower frequencies, since a low number of cycles fits in the investigated 300 ms time windows. While the interpretations of connections at specific frequencies remain unclear, the majority of our findings are consistent with the assumption that shorter connections utilize relatively higher frequencies (von Stein and Sarnthein, 2000).

To conclude, we found that core regions of the semantic control network engage differently during semantic cognition: While both regions increase their oscillatory power based on the presence of semantic goals, IFG constrains activation in pMTG. pMTG additionally receives early bottom-up input from visual cortex, suggesting that known semantic goals enable a rapid retrieval of task-relevant features. Our findings indicate that IFG *controls* goal-dependent retrieval, while pMTG *integrates* top-down semantic control with bottom-up input.

## Notes

### Competing Interest Statement

The authors have declared no competing interest.

### Summary of Updates

The main change is an improvement of the Granger Causality analysis by now using multivariate conditional Granger Causality.

### Funding

- European Commission: 771863
- Agencia Estatal de Investigación: CEX2020-001010/AEI/10.13039/501100011033

## References

1. Anwyl-Irvine AL, Massonnié J, Flitton A, Kirkham N, Evershed JK (2020) Gorilla in our midst: An online behavioral experiment builder. Behav Res 52:388–407.

2. Chiou R, Humphreys GF, Jung J, Lambon Ralph MA (2018) Controlled semantic cognition relies upon dynamic and flexible interactions between the executive ‘semantic control’ and hub-and-spoke ‘semantic representation’ systems. Cortex 103:100–116.

3. Davey J, Thompson HE, Hallam G, Karapanagiotidis T, Murphy C, De Caso I, Krieger-Redwood K, Bernhardt BC, Smallwood J, Jefferies E (2016) Exploring the role of the posterior middle temporal gyrus in semantic cognition: Integration of anterior temporal lobe with executive processes. NeuroImage 137:165–177.

4. Deniz F, Tseng C, Wehbe L, Tour TD la, Gallant JL (2023) Semantic Representations during Language Comprehension Are Affected by Context. J Neurosci 43:3144–3158.

5. Dhamala M, Rangarajan G, Ding M (2008) Analyzing information flow in brain networks with nonparametric Granger causality. NeuroImage 41:354–362.

6. Dikker S, Assaneo MF, Gwilliams L, Wang L, Kösem A (2020) Magnetoencephalography and Language. Neuroimaging Clin N Am 30:229–238.

7. Eisenhauer S, Fiebach CJ, Gagl B (2019) Context-Based Facilitation in Visual Word Recognition: Evidence for Visual and Lexical But Not Pre-Lexical Contributions. eNeuro 6:ENEURO.0321-18.2019.

8. Eisenhauer S, Gagl B, Fiebach CJ (2022) Predictive pre-activation of orthographic and lexical-semantic representations facilitates visual word recognition. Psychophysiology 59:e13970.

9. Fruchter J, Linzen T, Westerlund M, Marantz A (2015) Lexical preactivation in basic linguistic phrases. Journal of Cognitive Neuroscience 27:1912–1935.

10. Gao Z, Zheng L, Chiou R, Gouws A, Krieger-Redwood K, Wang X, Varga D, Ralph MAL, Smallwood J, Jefferies E (2021) Distinct and common neural coding of semantic and non-semantic control demands. NeuroImage 236:118230.

11. Geweke J (1982) Measurement of Linear Dependence and Feedback between Multiple Time Series. Journal of the American Statistical Association 77:304–313.

12. Giallanza T, Campbell D, Cohen JD, Rogers TT (2024) An integrated model of semantics and control. Psychological Review: Advance online publication.

13. Granger CWJ (1969) Investigating Causal Relations by Econometric Models and Cross-spectral Methods. Econometrica 37:424–438.

14. Grindrod CM, Bilenko NY, Myers EB, Blumstein SE (2008) The role of the left inferior frontal gyrus in implicit semantic competition and selection: An event-related fMRI study. Brain Research 1229:167–178.

15. Hallam GP, Whitney C, Hymers M, Gouws AD, Jefferies E (2016) Charting the effects of TMS with fMRI: Modulation of cortical recruitment within the distributed network supporting semantic control. Neuropsychologia 93:40–52.

16. Helbling S (2015) Advances in MEG methods and their applications to investigate auditory perception. PhD Thesis, Frankfurt, Goethe University.

17. Hohenstein S, Kliegl R (2015) remef: Remove partial effects. R package version 1.

18. Hrkać M, Wurm MF, Kühn AB, Schubotz RI (2015) Objects Mediate Goal Integration in Ventrolateral Prefrontal Cortex during Action Observation. PLOS ONE 10:e0134316.

19. Jackson RL (2021) The neural correlates of semantic control revisited. NeuroImage 224:117444.

20. Jackson RL, Hoffman P, Pobric G, Ralph MAL (2016) The Semantic Network at Work and Rest: Differential Connectivity of Anterior Temporal Lobe Subregions. J Neurosci 36:1490–1501.

21. Jackson RL, Rogers TT, Lambon Ralph MA (2021) Reverse-engineering the cortical architecture for controlled semantic cognition. Nat Hum Behav 5:774–786.

22. Jefferies E (2013) The neural basis of semantic cognition: Converging evidence from neuropsychology, neuroimaging and TMS. Cortex 49:611–625.

23. Krieger-Redwood K, Steward A, Gao Z, Wang X, Halai A, Smallwood J, Jefferies E (2023) Creativity in verbal associations is linked to semantic control. Cerebral Cortex 33:5135–5147.

24. Kuznetsova A, Brockhoff PB, Christensen RHB (2017) lmerTest Package: Tests in Linear Mixed Effects Models. Journal of Statistical Software 82:1–26.

25. Lau EF, Phillips C, Poeppel D (2008) A cortical network for semantics: (de)constructing the N400. Nature Reviews Neuroscience 9:920–933.

26. Mandera P, Keuleers E, Brysbaert M (2017) Explaining human performance in psycholinguistic tasks with models of semantic similarity based on prediction and counting: A review and empirical validation. Journal of Memory and Language 92:57–78.

27. Maris E, Oostenveld R (2007) Nonparametric statistical testing of EEG– and MEG-data. Journal of Neuroscience Methods 164:177–190.

28. McNab F, Rippon G, Hillebrand A, Singh KD, Swithenby SJ (2007) Semantic and phonological task-set priming and stimulus processing investigated using magnetoencephalography (MEG). Neuropsychologia 45:1041–1054.

29. Morawetz C, Bode S, Baudewig J, Jacobs AM, Heekeren HR (2016) Neural representation of emotion regulation goals. Human Brain Mapping 37:600–620.

30. Nolte G (2003) The magnetic lead field theorem in the quasi-static approximation and its use for magnetoencephalography forward calculation in realistic volume conductors. Phys Med Biol 48:3637–3652.

31. Noonan KA, Jefferies E, Visser M, Lambon Ralph MA (2013) Going beyond inferior prefrontal involvement in semantic control: evidence for the additional contribution of dorsal angular gyrus and posterior middle temporal cortex. J Cogn Neurosci 25:1824–1850.

32. Oostenveld R, Fries P, Maris E, Schoffelen J-M (2011) FieldTrip: Open source software for advanced analysis of MEG, EEG, and invasive electrophysiological data. Comput Intell Neurosci 2011:156869.

33. Peirce J, Gray JR, Simpson S, MacAskill M, Höchenberger R, Sogo H, Kastman E, Lindeløv JK (2019) PsychoPy2: Experiments in behavior made easy. Behav Res 51:195–203.

34. Prystauka Y, Lewis AG (2019) The power of neural oscillations to inform sentence comprehension: A linguistic perspective. Language and Linguistics Compass 13:e12347.

35. R Development Core Team (2008) R: a language and environment for statistical computing. Vienna, Austria: R Foundation for Statistical Computing.

36. Rahimi S, Farahibozorg S-R, Jackson R, Hauk O (2022) Task modulation of spatiotemporal dynamics in semantic brain networks: An EEG/MEG study. Neuroimage 246:118768.

37. Ralph MAL, Jefferies E, Patterson K, Rogers TT (2017) The neural and computational bases of semantic cognition. Nat Rev Neurosci 18:42–55.

38. Sassenhagen J, Draschkow D (2019). Cluster-based permutation tests of MEG/EEG data do not establish significance of effect latency or location. Psychophysiology 56:e13335.

39. Schaum M, Pinzuti E, Sebastian A, Lieb K, Fries P, Mobascher A, Jung P, Wibral M, Tüscher O (2021) Right inferior frontal gyrus implements motor inhibitory control via beta-band oscillations in humans Little S, Ivry RB, Little S, eds. eLife 10:e61679.

40. Schurz M, Kronbichler M, Crone J, Richlan F, Klackl J, Wimmer H (2014) Top-down and bottom-up influences on the left ventral occipito-temporal cortex during visual word recognition: An analysis of effective connectivity. Human Brain Mapping 35:1668–1680.

41. Schwartz MF, Kimberg DY, Walker GM, Brecher A, Faseyitan OK, Dell GS, Mirman D, Coslett HB (2011) Neuroanatomical dissociation for taxonomic and thematic knowledge in the human brain. Proceedings of the National Academy of Sciences 108:8520–8524.

42. Snyder HR, Banich MT, Munakata Y (2011) Choosing Our Words: Retrieval and Selection Processes Recruit Shared Neural Substrates in Left Ventrolateral Prefrontal Cortex. Journal of Cognitive Neuroscience 23:3470–3482.

43. Teige C, Cornelissen PL, Mollo G, Gonzalez Alam TR del J, McCarty K, Smallwood J, Jefferies E (2019) Dissociations in semantic cognition: Oscillatory evidence for opposing effects of semantic control and type of semantic relation in anterior and posterior temporal cortex. Cortex 120:308–325.

44. Teige C, Mollo G, Millman R, Savill N, Smallwood J, Cornelissen PL, Jefferies E (2018) Dynamic semantic cognition: Characterising coherent and controlled conceptual retrieval through time using magnetoencephalography and chronometric transcranial magnetic stimulation. Cortex 103:329–349.

45. Turken AU, Dronkers NF (2011) The Neural Architecture of the Language Comprehension Network: Converging Evidence from Lesion and Connectivity Analyses. Front Syst Neurosci 5:1.

46. Van Casteren M, Davis MH (2007) Match: A program to assist in matching the conditions of factorial experiments. Behavior Research Methods 39:973–978.

47. Van Veen BD, van Drongelen W, Yuchtman M, Suzuki A (1997) Localization of brain electrical activity via linearly constrained minimum variance spatial filtering. IEEE Trans Biomed Eng 44:867–880.

48. Voeten CC (2025) buildmer: Stepwise Elimination and Term Reordering for Mixed-Effects Regression. R package version 2.12, https://gitlab.com/cvoeten/buildmer.

49. von Stein A, Sarnthein J (2000) Different frequencies for different scales of cortical integration: from local gamma to long range alpha/theta synchronization. International Journal of Psychophysiology 38:301–313.

50. Wang X, Gao Z, Smallwood J, Jefferies E (2021) Both Default and Multiple-Demand Regions Represent Semantic Goal Information. J Neurosci 41:3679–3691.

51. Wawrzyniak M, Hoffstaedter F, Klingbeil J, Stockert A, Wrede K, Hartwigsen G, Eickhoff SB, Classen J, Saur D (2017) Fronto-temporal interactions are functionally relevant for semantic control in language processing. PLOS ONE 12:e0177753.

52. Woodhead ZVJ, Barnes GR, Penny W, Moran R, Teki S, Price CJ, Leff AP (2014) Reading front to back: MEG evidence for early feedback effects during word recognition. Cereb Cortex 24:817–825.

53. Wurm MF, Hrkać M, Morikawa Y, Schubotz RI (2014) Predicting goals in action episodes attenuates BOLD response in inferior frontal and occipitotemporal cortex. Behavioural Brain Research 274:108–117.

54. Xu Z, Shen B, Taji W, Sun P, Naya Y (2020) Convergence of distinct functional networks supporting naming and semantic recognition in the left inferior frontal gyrus. Human Brain Mapping 41:2389–2405.

55. Yue Q, Zhang L, Xu G, Shu H, Li P (2013) Task-modulated activation and functional connectivity of the temporal and frontal areas during speech comprehension. Neuroscience 237:87–95.

56. Zhang M, Varga D, Wang X, Krieger-Redwood K, Gouws A, Smallwood J, Jefferies E (2021) Knowing what you need to know in advance: The neural processes underpinning flexible semantic retrieval of thematic and taxonomic relations. NeuroImage 224:117405.

57. Zhen Z, Fang H, Liu J (2013) The Hierarchical Brain Network for Face Recognition. PLOS ONE 8:e59886.

