## Supplemental Material for "Controlled retrieval relies on directed interactions between semantic control regions and visual cortex: MEG evidence from oscillatory dynamics"

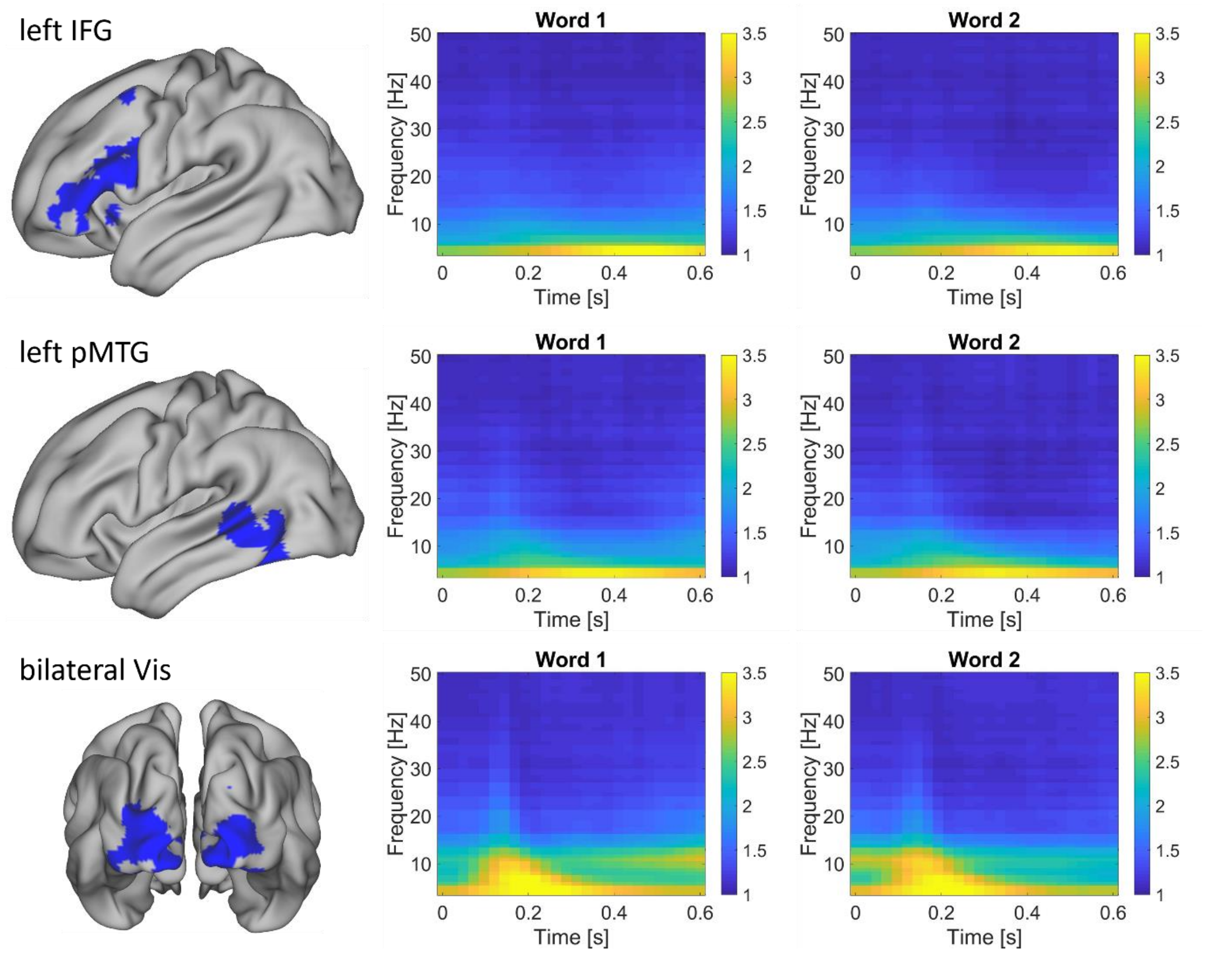


**Figure S1.** Brian regions of interest (left) and time-frequency representations of oscillatory power (middle, right), relative to the pre-cue baseline, in the left IFG (top), left pMTG (middle), and bilateral visual cortex (bottom) during the 600 ms presentation of word 1 and 2, averaged across knowledge conditions.

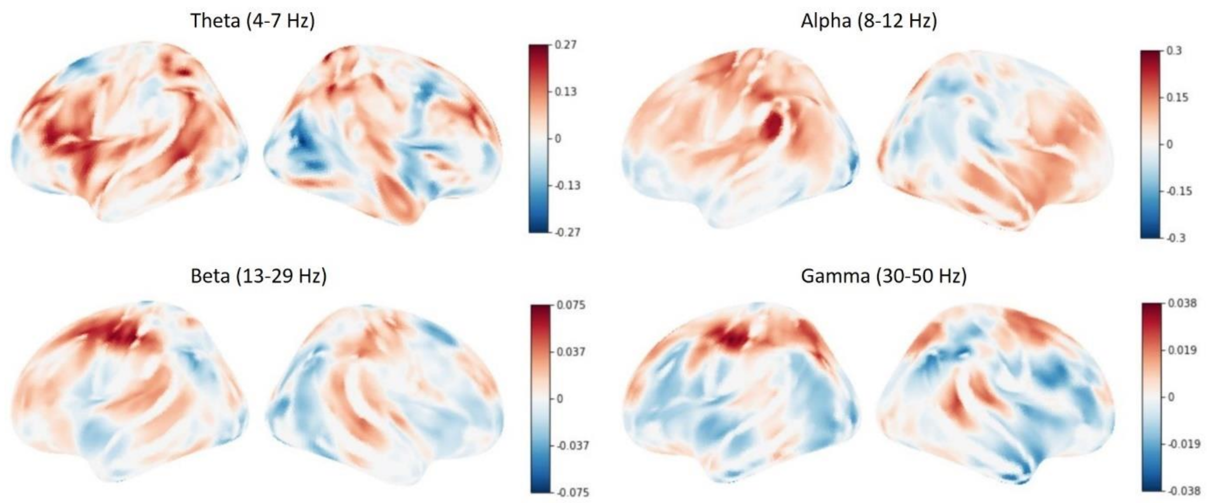


**Figure S2.** Whole-brain maps of oscillatory power, relative to the pre-cue baseline, for the contrast of known over unknown trials, averaged across the 600 ms during the presentation of word 1 and word 2, in theta, alpha, beta and gamma frequency ranges. Note that local peaks for known over unknown trials in theta power partly overlap with ROIs in the left inferior frontal and the left posterior middle temporal gyrus, and local peaks for unknown over known trials in alpha power partly overlap with the ROI in bilateral visual cortex.

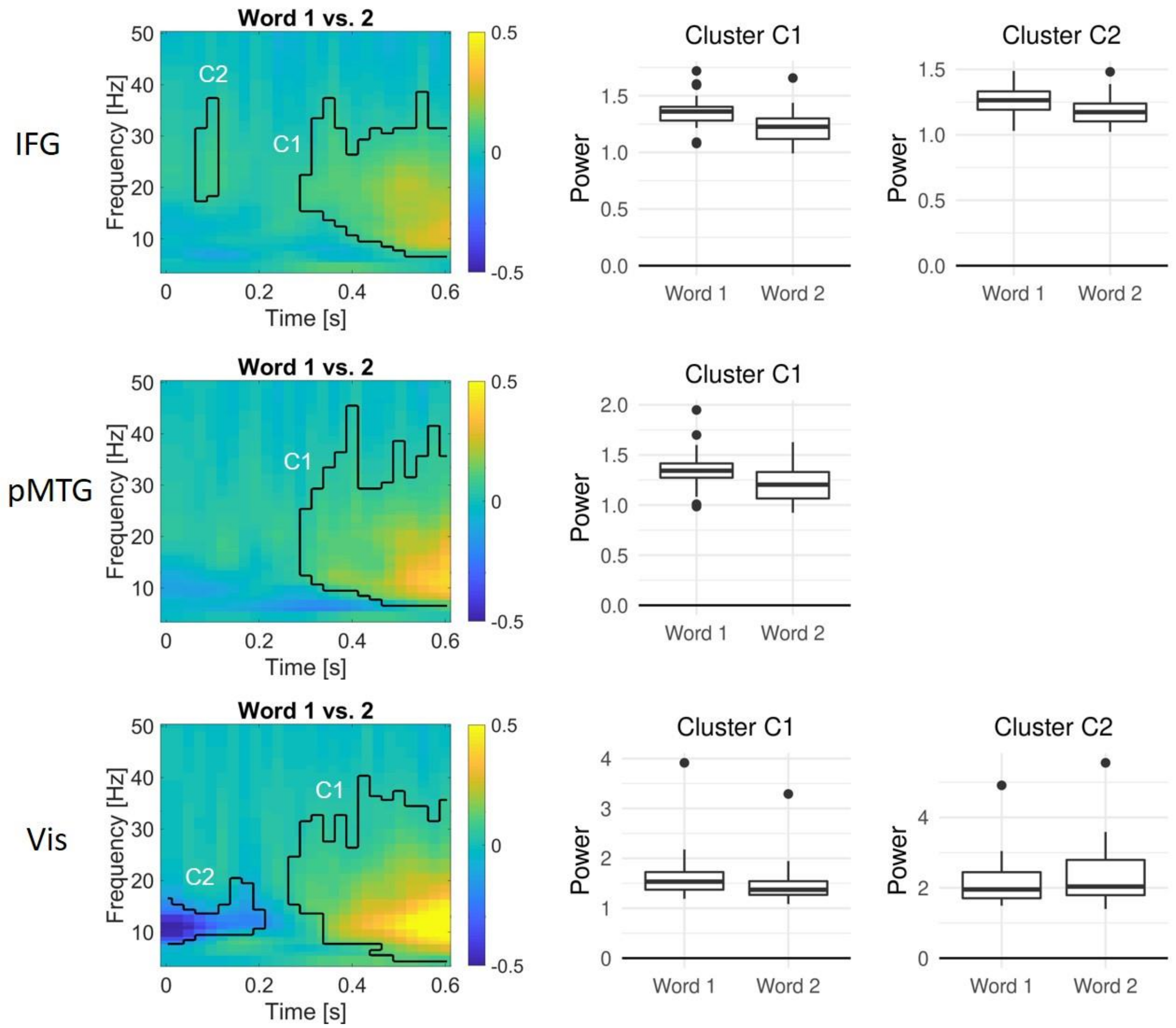


**Figure S3.** Oscillatory dynamics of the main effect of word order in left IFG (top), left pMTG (middle), and bilateral visual cortex (bottom), locked to word onset. Time-frequency plots show the difference in power (relative to the pre-cue baseline) between word 1 and 2, averaged across knowledge conditions, in significant clusters. Boxplots show the oscillatory power in each significant cluster for word 1 and 2 across participants.

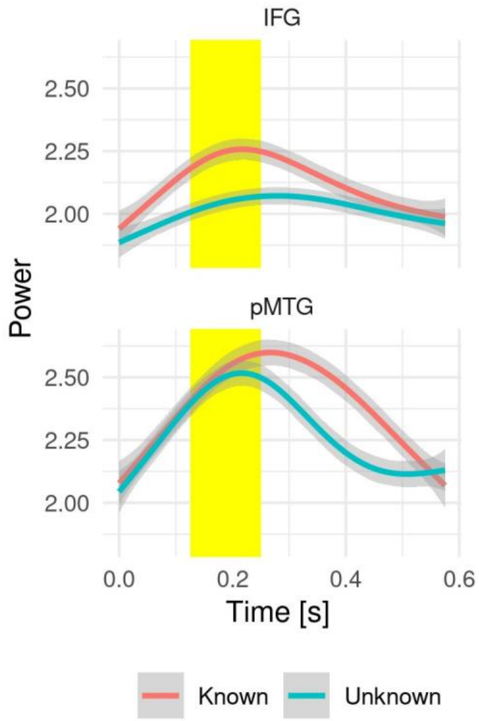


**Figure S4.** A follow-up analysis focused on the frequency where the known over unknown effect on oscillatory power in IFG and pMTG was maximal (7 and 6 Hz, respectively, Fig. 4) revealed main effects of knowledge from 125 to 250 ms (yellow shading, $p < 0.05$, not corrected for multiple comparisons across time points), but no significant interactions with ROI (IFG vs. pMTG). Grey shading represents 95 % confidence intervals. Trial-level data were analysed using linear mixed models including fixed effects of task knowledge (known vs. unknown goal, averaged across words 1 and 2), ROI (IFG vs. pMTG) and their interaction, as well as random intercepts of participant and item, and random slopes of all fixed effects for participants and items. Since these maximal models did not converge, the buildmer package (Voeten, 2025) was used to identify the maximal random slopes structure that converged at each time point (singular fit warnings were accepted since they could not be avoided for some models).
